# Meiotic Cells Escape Prolonged Spindle Checkpoint Activity Through Premature Silencing and Slippage

**DOI:** 10.1101/2023.01.02.522494

**Authors:** Anne MacKenzie, Victoria Vicory, Soni Lacefield

## Abstract

To prevent chromosome mis-segregation, a surveillance mechanism known as the spindle checkpoint delays the cell cycle if kinetochores are not attached to spindle microtubules, allowing the cell additional time to correct improper attachments. During spindle checkpoint activation, checkpoint proteins bind the unattached kinetochore and send a diffusible signal to inhibit the anaphase promoting complex/cyclosome (APC/C). Previous work has shown that mitotic cells with depolymerized microtubules can escape prolonged spindle checkpoint activation in a process called mitotic slippage. During slippage, spindle checkpoint proteins bind unattached kinetochores, but the cells cannot maintain the checkpoint arrest. We asked if meiotic cells had as robust of a spindle checkpoint response as mitotic cells and whether they also undergo slippage after prolonged spindle checkpoint activity. We performed a direct comparison between mitotic and meiotic budding yeast cells that signal the spindle checkpoint due to a lack of either kinetochore-microtubule attachments or due to a loss of tension-bearing attachments. We find that the spindle checkpoint is not as robust in meiosis I or meiosis II compared to mitosis, overcoming a checkpoint arrest approximately 150 minutes earlier in meiosis. In addition, cells in meiosis I escape spindle checkpoint signaling using two mechanisms, silencing the checkpoint at the kinetochore and through slippage. We propose that meiotic cells undertake developmentally-regulated mechanisms to prevent persistent spindle checkpoint activity to ensure the production of gametes.

**AUTHOR SUMMARY:** Mitosis and meiosis are the two major types of cell divisions. Mitosis gives rise to genetically identical daughter cells, while meiosis is a reductional division that gives rise to gametes. Cell cycle checkpoints are highly regulated surveillance mechanisms that prevent cell cycle progression when circumstances are unfavorable. The spindle checkpoint promotes faithful chromosome segregation to safeguard against aneuploidy, in which cells have too many or too few chromosomes. The spindle checkpoint is activated at the kinetochore and then diffuses to inhibit cell cycle progression. Although the checkpoint is active in both mitosis and meiosis, most studies involving checkpoint regulation have been performed in mitosis. By activating the spindle checkpoint in both mitosis and meiosis in budding yeast, we show that cells in meiosis elicit a less persistent checkpoint signal compared to cells in mitosis. Further, we show that cells use distinct mechanisms to escape the checkpoint in mitosis and meiosis I. While cells in mitosis and meiosis II undergo anaphase onset while retaining checkpoint proteins at the kinetochore, cells in meiosis I prematurely lose checkpoint protein localization at the kinetochore. If the mechanism to remove the checkpoint components from the kinetochore is disrupted, meiosis I cells can still escape checkpoint activity. Together, these results highlight that cell cycle checkpoints are differentially regulated during meiosis to avoid long delays and to allow gametogenesis.

## INTRODUCTION

A conserved surveillance mechanism known as the spindle checkpoint delays chromosome segregation in both mitosis and meiosis if kinetochores are not attached to spindle microtubules [1]. The delay provides cells additional time to make proper attachments prior to the onset of chromosome segregation. Although the spindle checkpoint reduces chromosome mis-segregation, segregation errors can still occur and cause aneuploidy, in which cells have missing or extra chromosomes. Aneuploidy can be detrimental to normal cellular function and can contribute to human disease [2]. For example, aneuploidy is a hallmark of cancer and leads to genetic instability and cancer progression. Aneuploidy during meiosis contributes to trisomy conditions, miscarriage, and infertility. Therefore, understanding how the spindle checkpoint normally functions and how it can sometimes fail to prevent chromosome mis-segregation is important for understanding the etiology of these human diseases.

The spindle checkpoint produces a diffusible signal at an unattached kinetochore that ultimately inhibits the anaphase promoting complex/ cyclosome (APC/C), a ubiquitin ligase that targets substrates for ubiquitination and proteasomal degradation for anaphase onset [1]. At an unattached kinetochore, the spindle checkpoint kinase Mps1 phosphorylates the kinetochore protein Knl1^Spc105^ for the Bub3-Bub1 complex to bind [3–7]. Bub1 is phosphorylated by Mps1 and binds Mad1 to create a platform that recruits other checkpoint components, which ultimately assemble into the mitotic checkpoint complex (MCC) composed of Bub3, Mad2, Mad3/BubR1, and Cdc20 [8–14] [15, 16]. The MCC serves as the diffusible signal that inhibits the APC/C [17–19]. The spindle checkpoint is turned off once proper kinetochore-microtubule attachments are made in a process called spindle checkpoint silencing. Protein phosphatase I (PP1) binds and dephosphorylates Knl1^Spc105^ to remove Bub3, preventing production of the MCC [3, 20–22]. Furthermore, the MCC is disassembled allowing the APC/C to become active for anaphase onset by targeting securin and cyclin B for ubiquitination and subsequent degradation[1].

Cells can escape the spindle checkpoint after a prolonged delay [23]. For example, experimental conditions that completely disrupt kinetochore-microtubule attachments, such as with the addition of the microtubule depolymerizing drug nocodazole, cause an arrest for several hours. Interestingly, cells prevent a permanent arrest by escaping the spindle checkpoint in a process known as mitotic slippage or adaptation [24–27]. Mitotic slippage occurs in budding yeast and animal cells despite the presence of spindle checkpoint components bound to the kinetochore [24, 27]. During mitotic slippage, securin and cyclin B are degraded for anaphase onset, due to activation of the APC/C, MCC disassembly, cyclin B turnover, or Cdk1 inhibition [24, 28, 29]. One mechanism that allows mitotic slippage in budding yeast is through PP1. Instead of PP1 targeting the kinetochore, as occurs during spindle checkpoint silencing, PP1 targets the MCC component Mad3 during mitotic slippage [29]. The dephosphorylation of Mad3 destabilizes the MCC, allowing APC/C activation for anaphase onset.

Most experiments on spindle checkpoint activity and slippage have been performed in mitosis. However, meiosis poses additional challenges to chromosome segregation that may require differences in spindle checkpoint regulation. Meiosis consists of two divisions in which homologous chromosomes segregate during meiosis I and sister chromatids segregate during meiosis II. Setting up the specialized segregation pattern in meiosis I requires that homologous chromosomes pair and undergo crossover recombination to form linkages between the chromosomes [30]. The two sister chromatid kinetochores are held together to form one microtubule binding site so that they co-orient. This allows a connection of homologous chromosome kinetochores to microtubules emanating from opposite spindle poles for biorientation. The poleward spindle forces are resisted by the crossover and the cohesins along the chromosome arms until the cohesins are cleaved in anaphase I for chromosome segregation [31–34]. The meiosis II chromosome segregation pattern is similar to that of mitosis, in that sister chromatid kinetochores attach to microtubules emanating from opposite spindle poles [30]. The spindle forces are resisted by cohesins in metaphase II until they are cleaved and sister chromatids separate. Whether these differences in chromosome segregation and cell cycle regulation affect spindle checkpoint activity is not known.

We performed a direct comparison of spindle checkpoint strength between mitosis and meiosis in budding yeast. We monitored the length of the checkpoint delay signaled through either a lack of attachment or tension-bearing kinetochore-microtubule attachments. The lack of tension-bearing attachments signals the spindle checkpoint through the activity of Aurora B kinase, which phosphorylates kinetochore proteins to release microtubules, creating unattached kinetochores [30]. Whether there is a difference in checkpoint strength between a lack of attachment versus lack of tension is an important question because Aurora B kinase localizes to both the kinetochore and microtubules and is involved in maintenance of spindle checkpoint activity [35–40]. Whether the pool of Aurora B kinase on the microtubules affects checkpoint activity and whether there are differences between mitosis and meiosis in the signaling is currently unknown. To this end, we found that cells escaped spindle checkpoint signaling significantly faster in both meiosis I and meiosis II when compared to mitosis, signaled through either addition of a microtubule-depolymerizing drug or a genetic background that caused a lack of intra-kinetochore tension. In contrast to mitosis and meiosis II, in which cells undergo mitotic slippage, cells in meiosis I can escape the spindle checkpoint through two mechanisms, slippage and silencing. Our data in budding yeast support the model that meiosis has a developmentally controlled regulation to turn off spindle checkpoint activity to ensure completion of meiosis for gamete formation.

## RESULTS

### The spindle checkpoint is less persistent in meiosis compared to mitosis

To investigate differences in spindle checkpoint strength between mitosis and meiosis, we used live-cell fluorescence microscopy to monitor the duration of the spindle checkpoint delay. The cells expressed the spindle pole body (SPB) component Spc42 tagged with mCherry and a biosensor for the enzyme separase, which cleaves cohesin at the metaphase-to-anaphase transition [41]. The biosensor consists of a chromosomal locus tagged with a fluorescent focus that disappears upon separase activation. To this end, a LacO array is integrated into a chromosome and the cells express a GFP-tagged Lac repressor engineered with an Scc1 cleavage site (GFP-Scc1-LacI) for mitosis or a Rec8 cleavage site for meiosis (GFP-Rec8-LacI). When separase is active, the cohesin subunit Scc1 or Rec8 will be cleaved and a GFP focus will no longer be visible, although the nucleus will still have a GFP haze (Figure 1A-B). The advantage of using the separase biosensor is that we can then monitor anaphase onset in conditions or mutant backgrounds that may have depolymerized microtubules or spindle elongation prior to cohesin cleavage (Figure 1C-D).

**Figure 1.**
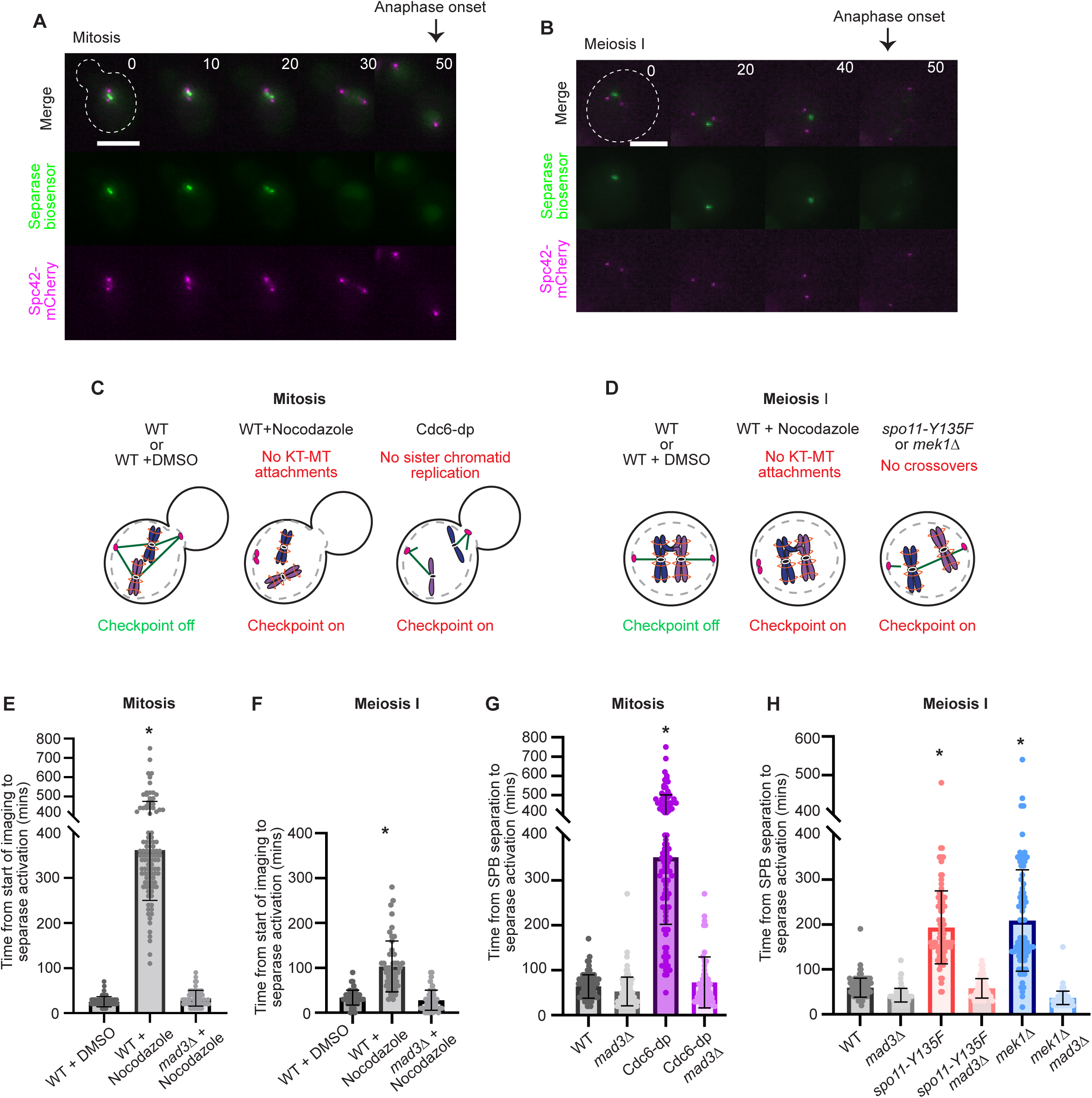
The spindle checkpoint is less persistent in meiosis compared to mitosis. (A-B) Representative time lapse images of a cell harboring Spc42-mCherry and the separase biosensor in mitosis (A) and meiosis I (B). Time from SPB separation to separase activation. Time 0 indicates SPB separation. Scale bar = 5 µm. (C-D) Cartoons depicting activation of the spindle checkpoint through loss of kinetochore (KT) tension and loss of kinetochore-microtubule (KT-MT) attachments in mitosis (C) and meiosis I (D) Wildtype = WT. E) Graph depicting mean time from start of movie to anaphase onset in mitosis following treatment with DMSO or nocodazole. Asterisk indicates statistically significant difference between DMSO-treated and nocodazole-treated cells (p <0.05, Mann-Whitney test) and error bars show standard deviation (SD) n≥ 50 cells per genotype. F) Graph depicting mean time from start of movie to anaphase onset in meiosis following treatment with DMSO or nocodazole. (G-H) Graph showing the mean time from SPB separation to anaphase onset in mitosis (G) and meiosis I (H). Asterisks indicate statistically significant difference between wildtype and mutants (p < 0.05, Mann-Whitney test). Error bars show SD. n≥100 cells per genotype.

We first measured the duration of a spindle checkpoint delay in a condition in which kinetochore-microtubule attachments are disrupted due to treatment with the microtubule-depolymerizing drug nocodazole (Figure 1C-D). In mitosis, we released haploid cells from an alpha-factor-induced G1 arrest and treated cells with 15 µg/mL of nocodazole after we observed separation of SPBs. We then started the movie and measured the duration from the start of the movie until loss of the GFP focus, which represents the time of separase cleavage of cohesin. We observed a metaphase duration of 348 ± 93 minutes when cells were treated with nocodazole, which is a delay of 323 minutes compared to cells treated with the solvent control (DMSO) (average ± SD Figure 1E). As previously shown for mitotic cells, we observed cell-to-cell variability in the duration of the arrest [27]. This delay is dependent on the spindle checkpoint because *mad3Δ* cells treated with nocodazole progressed into anaphase with a similar duration as the DMSO control cells.

To assess the duration of a checkpoint delay in meiosis I, we treated cells with 30 µg/mL of nocodazole after release from a prophase I arrest using the *GAL-NDT80* arrest and release system [42, 43]. We treated cells with nocodazole 80-minutes after release from prophase I, which is when we observed SPB separation. We specifically waited to add the nocodazole until after prophase I because microtubule perturbation earlier causes a G2 arrest [44]. In the presence of nocodazole, cells had a metaphase I duration of 103 ± 56 minutes, which is 69 minutes longer than the control cells treated with DMSO alone (average ± SD; Figure 1F). The *mad3Δ* cells treated with nocodazole had a similar time of anaphase onset as the DMSO control, suggesting that the delay was spindle checkpoint dependent. We conclude that the spindle checkpoint is more persistent in mitosis than in meiosis when kinetochore-microtubule attachments are disrupted.

Chromosome segregation errors that cause single chromosome aneuploidies often arise from the misattachment of kinetochores and microtubules, such that the two sister chromatids in mitosis or the two homologous chromosomes in meiosis I are attached to the same pole. The monopolar attachment creates a lack of intra-kinetochore stretch, or tension [45]. This lack of tension can activate the Ipl1^Aurora^ ^B^ kinase to release inappropriate attachments, ultimately leading to spindle checkpoint activation [46–49]. We therefore wanted to compare the persistence of the spindle checkpoint delay between mitosis and meiosis I upon disruption of kinetochore tension. We engineered strains to create a condition in which kinetochores are attached to spindle microtubules but cannot biorient, creating a lack of tension (Figure 1C-D).

To prevent kinetochore tension in mitosis, we depleted the essential protein Cdc6, which is required for initiation of DNA replication (Piatti, 1995). Because only one microtubule binds per kinetochore in budding yeast, without DNA replication, the single sister chromatid will be unable to biorient, creating a lack of kinetochore tension. To deplete Cdc6, *CDC6* was placed under control of the *GAL1* galactose-inducible promoter and fused to a Ubi degron (Cdc6-dp) [50]. When cells are grown in the presence of glucose, *CDC6* expression is repressed, and any remaining Cdc6 is degraded. We assessed the duration of the spindle checkpoint delay in diploid Cdc6-dp cells by measuring the time from SPB separation to separase activation, as assessed by the loss of the GFP focus. Cdc6-dp cells were in metaphase for a duration of 352 ± 148 minutes, which is a delay of 295 minutes when compared to wildtype cells (average ± SD; Figure 1G). The observed delay in Cdc6-dp cells was due to spindle checkpoint activation because Cdc6-dp *mad3Δ* cells progressed through metaphase with similar timings as the wildtype control cells. There was also cell-to-cell variability in the duration of the checkpoint delay in Cdc6-dp cells, similar to nocodazole treated cells.

We note that the timings from the lack of tension experiments cannot be directly compared to the timings from the nocodazole experiments. In the nocodazole experiments, we did not measure the full duration of metaphase because time-lapse imaging was started after the addition of nocodazole, to prevent the drug from interfering with SPB separation (see Materials and Methods). In contrast, we measured the time from SPB separation to anaphase onset in the Cdc6-dp experiments. However, the results show that both nocodazole and Cdc6-dp cause an extended spindle checkpoint delay that cells can eventually escape.

To prevent intra-kinetochore tension during meiosis I, we disrupted proteins needed for crossover formation. Normally, crossovers, held in place by arm cohesin, link homologs together and resist the spindle forces when homologous chromosomes biorient, which creates intra-kinetochore tension [30]. We prevented crossover formation by using a catalytically inactive allele of *SPO11*, *spo11-Y135F*, which is a mutant form of the enzyme required to make double-stranded breaks for crossover formation [51]. Without crossovers, homologs are not linked, causing a lack of tension across the kinetochores. In these cells, chromosomes move to the spindle pole prematurely in an anaphase-like prometaphase I because they cannot resist spindle forces [52]. The *spo11-Y135F* cells exhibited a metaphase I duration of 194 ± 81 minutes, which is a delay of 140 minutes compared to the wildtype control cells (average ± SD; Fig 1H). The *spo11-Y135F mad3*Δ cells showed similar timings to the wildtype control cells, suggesting that the delay was spindle checkpoint dependent.

Like the mitotic studies, we cannot directly compare the nocodazole-treated cells with the *spo11-Y135F* cells. In the nocodazole experiment, we started the time-lapse imaging after addition of the nocodazole to cells in early metaphase I, just after SPB separation. In contrast, the *spo11-Y135F* cells were imaged throughout meiosis, providing the full duration of the metaphase I delay.

To determine if the duration of the spindle checkpoint delay is similar in other mutants with reduced crossovers, we deleted *MEK1*, which biases repair of DSBs to the homolog, allowing for crossover formation. In cells harboring *MEK1* mutants, crossover formation is reduced by ∼85%, so some homologs will retain crossovers [53–55]. The *mek1Δ* cells displayed a metaphase I delay of approximately 130 minutes with cell-to-cell variability, which were similar timings to those observed in *spo11-Y135F* cells (Figure 1H). Overall, our results demonstrate that the spindle checkpoint is not as persistent in meiosis as it is in mitosis, signaled through either a lack of kinetochore tension or a lack of attachment.

We wondered whether the shorter duration of the spindle checkpoint in meiosis was due to the size differences between budding yeast meiotic and mitotic cells. Mouse oocytes have a weaker spindle checkpoint, which is thought to be due to their larger size [56–63]. Similarly, checkpoint strength is thought to be at least partially dependent on cell size in the oocyte and developing *C. elegans* embryo [64–66]. To determine if a size difference could be causing a weaker spindle checkpoint in meiotic budding yeast cells, we measured the volume of budding yeast cells at the time point just before anaphase onset and found that *mek1Δ* cells in meiosis had a volume of 254 ± 46 fL, while Cdc6-dp cells had an average volume of 343 ± 89 fL (Figure S1). From this we conclude that cells arrested in mitosis have a larger average volume compared to cells arrested in meiosis, so it is not likely that the inherent weakness of the meiotic checkpoint is due to a larger cell volume.

### The meiotic spindle checkpoint prevents some chromosome mis-segregation

Our results showing that the meiotic spindle checkpoint delay is significantly shorter than the mitotic delay led us to question whether the short meiotic delay was sufficient to allow the correction of erroneous kinetochore-microtubule attachments. To address this question, we compared chromosome segregation fidelity between *mek1*Δ cells and *mek1*Δ *mad3*Δ cells, which lacked spindle checkpoint activity. We monitored the segregation of a single pair of homologs with LacO repeats near the centromere of chromosome IV in cells that expressed LacI-GFP under the control of a copper-inducible *CUP1* promoter [67]. Upon the addition of copper sulfate, LacI-GFP was expressed and bound the LacO repeats, which allowed visualization of each homolog as a single focus (Figure 2A). In wildtype and *mad3*Δ cells, the percent of chromosome IV mis-segregation was less than 2%. In contrast, 24% of *mek1*Δ and 38% of *mek1*Δ *mad3*Δ cells mis-segregated chromosome IV in anaphase I (Figure 2B). Thus, the meiotic spindle checkpoint prevented some chromosome mis-segregation in anaphase I, despite not having a robust delay.

**Figure 2.**
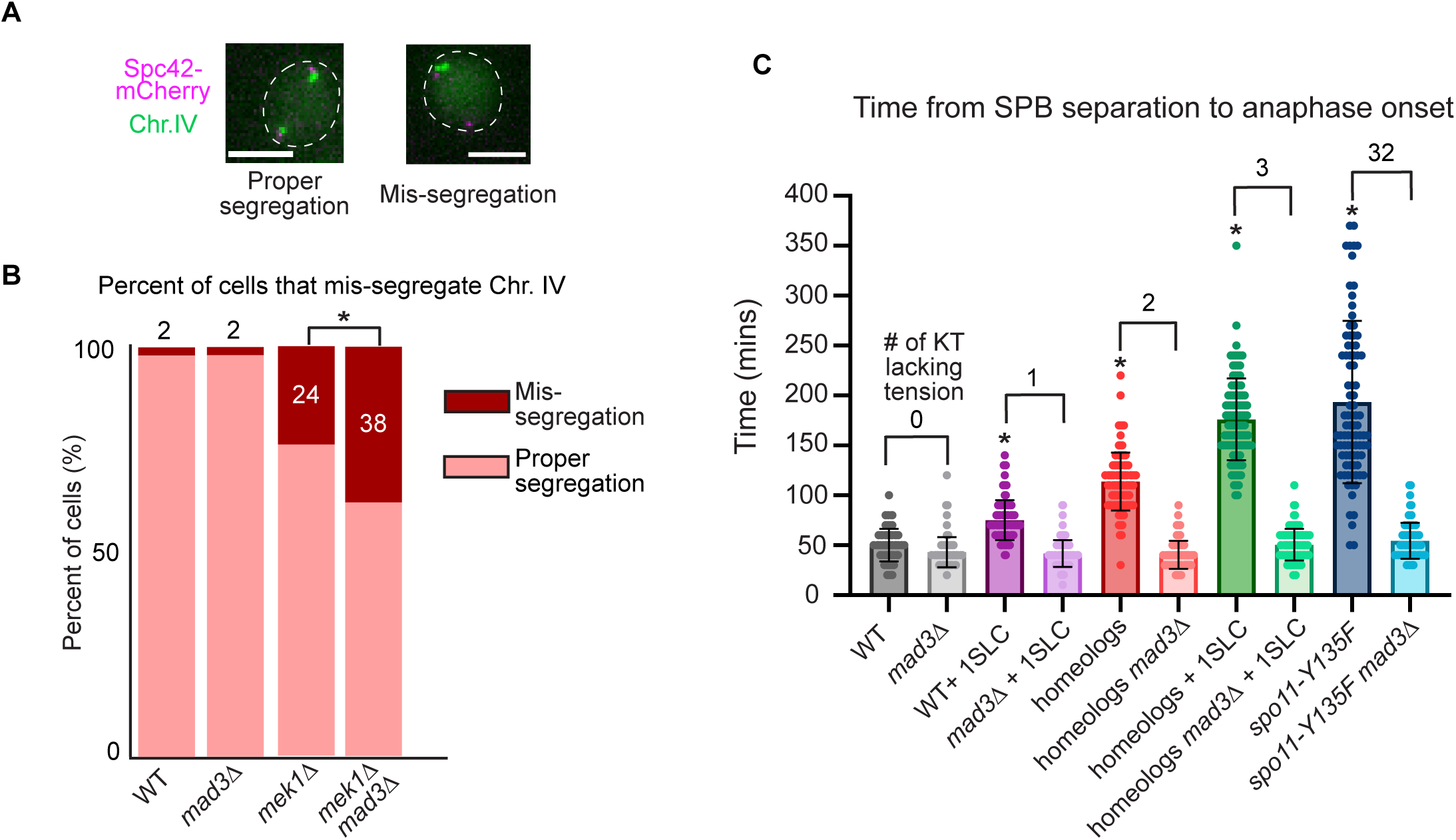
Spindle checkpoint activation reduces chromosome mis-segregation in meiosis I and is signaled to its full extent with three kinetochores that are not under tension. A) Representative images of a cell in which both homologs of chromosome IV are tagged with a GFP focus at anaphase I. Scale bar= 5 µm. B) Graph of percent mis-segregation of chromosome IV during meiosis I. Asterisk indicates a statistically significant difference from wildtype (Fisher’s Exact test). n≥ 100 cells per genotype. C) Graph of mean time from SPB separation to separase biosensor activation. Numbers above bars indicate the number of kinetochores lacking tension in each genotype. Asterisk represents statistically significant difference between wildtype and the indicated genotype (p <0.05, Mann-Whitney test), and error bars show SD. n≥100 cells per genotype. KT= kinetochore. SLC= short linear chromosome.

### The meiotic spindle checkpoint is signaled to its full extent when three kinetochores lack tension

Given the similarity in checkpoint duration between *spo11-Y135F* and *mek1Δ* cells in meiosis I, we questioned whether a single tensionless kinetochore could signal the full spindle checkpoint delay or if a threshold number of tensionless kinetochores was needed for the full delay. To this end, we engineered budding yeast strains with increasing numbers of kinetochores lacking tension and measured the time from SPB separation to separase activation. First, we introduced one short linear chromosome (SLC) into an otherwise wildtype yeast strain (WT + SLC). SLCs are not thought to form crossovers due to their short size and inability to hold the arm cohesins in place to stabilize the crossover [68–70]. But, the SLC has a centromere on which a kinetochore can build. The WT + SLC strain displayed a metaphase I duration of 75 ± 20 minutes, which is a checkpoint delay of 24 minutes compared to wildtype cells (average ± SD; Figure 2C). We engineered a strain in which two kinetochores lacked tension by introducing a single pair of homeologous chromosomes, which are chromosomes from two different yeast species that cannot form crossovers due to their sequence divergence [71, 72]. The strain harboring the homeologs displayed a checkpoint delay of 62 minutes compared to wildtype cells (Figure 2C). Next, we created a strain with three kinetochores that were not under tension, by adding a SLC to the strain harboring the homeologs (homeologs +1SLC). These cells underwent metaphase I after 176 ± 41 minutes, which is a checkpoint delay of 125 minutes, similar to that of *spo11-Y135F* cells (Figure 2C). Because *spo11-Y135F* cells do not have any of the 32 kinetochores under tension, we conclude that three tensionless kinetochores is the minimum number required for a full meiotic checkpoint delay.

### Mps1 and Ipl1 maintain the spindle checkpoint delay in cells with tensionless attachments

Because the spindle checkpoint delay was shorter in meiosis compared to mitosis, we next asked whether the Ipl1^Aurora^ ^B^ and Mps1 kinases were needed to maintain the meiotic spindle checkpoint delay. In meiosis, Ipl1^Aurora^ ^B^ releases improper attachments and Mps1 helps form force-generating attachments [73]. We used the anchor away technique to deplete Ipl1^Aurora^ ^B^ or Mps1 from the nucleus [74] (Figure S2A). We tagged endogenous *IPL1* or *MPS1* at its C-terminus with FRB in a strain with the ribosome protein *RPL13A* tagged with FKBP12. Because Rpl13A shuttles from the nucleus into the cytosol during ribosome biogenesis, the addition of rapamycin, which allows the stable interaction between FRB and FKBP12, depleted Ipl1-FRB or Mps1-FRB from the nucleus. We added rapamycin at 8 hours, a timepoint when many of the cells were in metaphase I, and then began live-cell imaging. We measured the time of anaphase I onset only in the cells that were in metaphase I at the start of imaging. We found that anchoring away of either Ipl1 or Mps1 in *mek1Δ* cells accelerated the duration of metaphase by 42 minutes and 53 minutes, respectively (Figure S2B). Therefore, both Ipl1^Aurora^ ^B^ and Mps1 maintained the spindle checkpoint delay in *mek1Δ* cells.

### Chromosomes undergo error correction attempts throughout the delay

The requirement of Ipl1^Aurora^ ^B^ for the spindle checkpoint delay in *mek1Δ* cells led us to question whether the relatively short checkpoint delay observed in *mek1Δ* cells was due to a decrease in Ipl1^Aurora^ ^B^ localization or activity over time. We first measured the level of Ipl1-3GFP that co-localized with the kinetochore protein Mtw1-mRuby2 at anaphase I onset. We found no significant difference in Ipl1-3GFP kinetochore levels in wildtype compared to *mek1Δ* cells (Figure S3).

To address the question of whether error correction activity decreased during the delay, we monitored the number of error correction attempts of one pair of homologs throughout a *mek1Δ* delay. We added a LacO array approximately 12kb from the centromere of both copies of chromosome IV. In the absence of Ipl1^Aurora^ ^B^ in meiosis, improper kinetochore-microtubule attachments are not corrected and both homologs stay attached to the same SPB [73, 75]. Therefore, to ask whether Ipl1 activity was attenuated during the meiotic spindle checkpoint delay, we counted the number of error correction attempts during a *mek1Δ* spindle checkpoint delay by following the detachment and reattachment of a single pair of homologs between SPBs, taking images every 5 minutes. We scored chromosomes as bioriented and non-bioriented and noted when a chromosome traversed between two SPBs. We considered any traversing from one pole to another or switching from non-bioriented to bioriented as an error correction attempt (Figure 3A).

**Figure 3.**
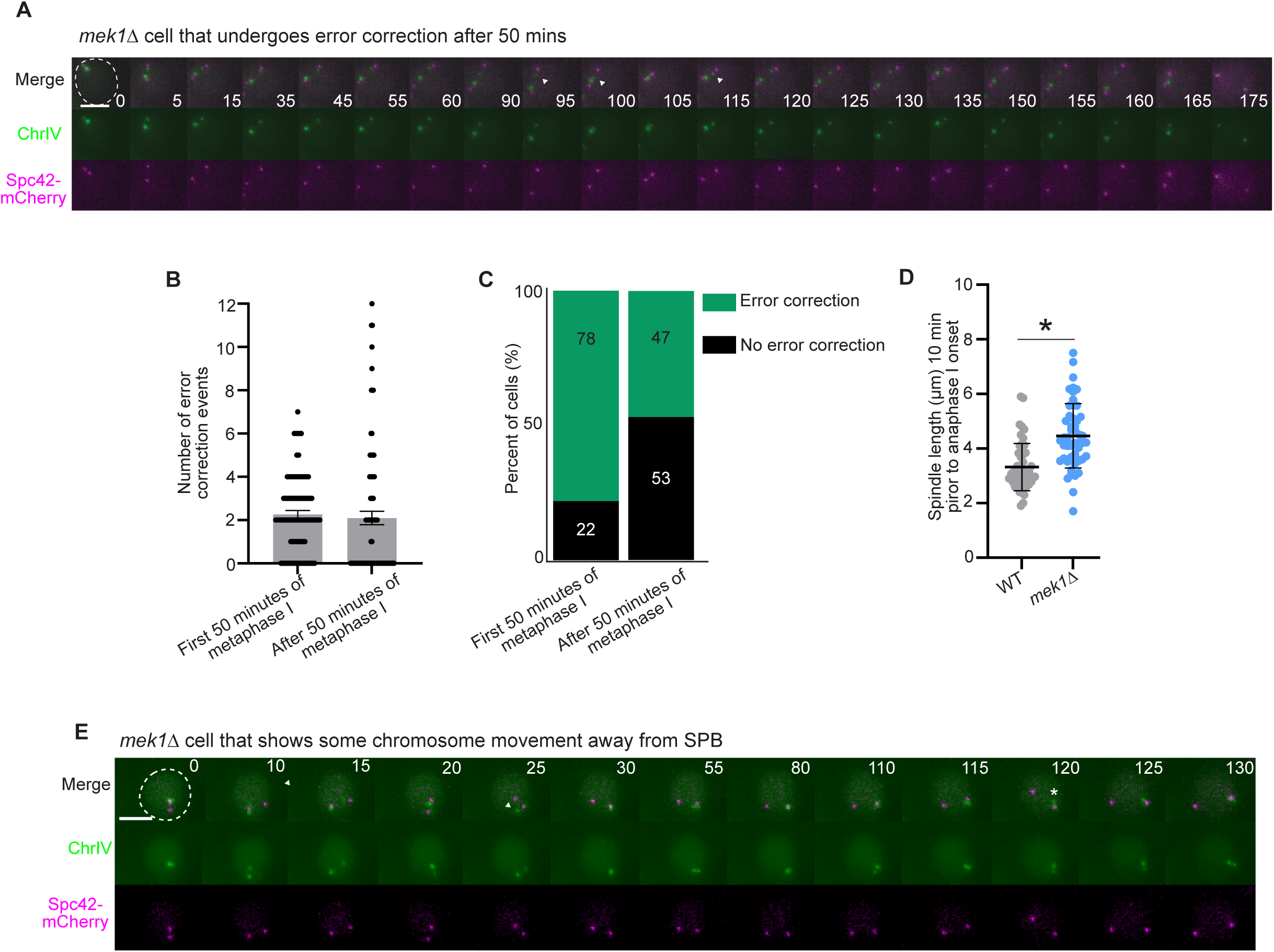
Chromosomes undergo error correction attempts throughout the spindle checkpoint delay in *mek1Δ* cells. A) Representative time-lapse images of a *mek1Δ* cell undergoing error correction after 50 minutes. Time 0 is the time at which SPBs separate. Arrowheads show movement of chromosome between SPBs. Scale bar=5 µm. B) Graph of individual error correction attempts per cell within the first 50 minutes and after the first 50 minutes of metaphase I. n≥ 100 cells. Error bars show SD. C) Percent of cells that underwent error correction in the first 50 minutes of metaphase I and after 50 minutes of metaphase I. n≥ 100 cells. D) Graph showing spindle length, measured 10 minutes prior to anaphase onset. n≥ 50 cells per genotype. Error bars show SD. Asterisk indicates a statistically significant difference between wildtype and *mek1Δ* (p < 0.05, Mann-Whitney test). E) Time-lapse images of a cell that does not undergo error correction after 50 minutes of metaphase I. Time 0 is the time at which SPBs separate. Arrowheads show chromosome movement between SPBs. Asterisk indicates chromosomes straying from SPB. Scale bar =5 µm.

We split our analysis into two categories, the number of events in the first 50 minutes of metaphase I and the number of error correction events after cells were delayed for 50 minutes. Because the time of anaphase onset was approximately 50 minutes in wildtype cells, we reasoned that the additional delay in *mek1Δ* cells was beyond 50 minutes. We found that 78% of *mek1Δ* cells underwent error correction events in the first 50 minutes of the delay, with an average of 2.3 ± 1.7 error correction attempts per cell (average ± standard deviation) (Figure 3B-C). After 50 minutes, 47% of *mek1Δ* cells still underwent error correction attempts, with a total average of 2.1 ± 3 error correction attempts. We hypothesized that more cells attempted error correction, but that the event did not hold up to our criteria. For example, the *mek1Δ* cells had a longer spindle length than wildtype cells prior to anaphase I onset, likely due to the prolonged spindle forces (3.3um ± 0.9 μm in wildtype compared to 4.5um ± 1.2 μm in *mek1Δ* cells; average ± SD; Figure 3D). The increased spindle length may have made it harder for kinetochores to attach to microtubules emanating from the opposite SPB, and instead, the released kinetochore reattached to the same SPB. We noticed that the LacI-GFP focus often drifted from the SPB, suggesting that the chromosome was released from the SPB, but may not have been able to traverse or biorient (Figure 3E). Therefore, because we observed error correction throughout the spindle checkpoint delay in *mek1Δ* cells, we conclude that premature attenuation of Ipl1 activity is not responsible for the short meiotic spindle checkpoint delay.

### The spindle checkpoint is silenced prematurely in meiosis I but not in mitosis or meiosis II

We wondered why the spindle checkpoint delay was shorter in meiosis compared to mitosis. To address this question, we asked how cells escaped the spindle checkpoint. Progression past the spindle checkpoint occurs through various mechanisms, which can be classified into two major categories. First, although checkpoint silencing normally occurs to turn off the checkpoint after chromosome biorientation in mitosis, we hypothesized that checkpoint silencing could inappropriately occur during a prolonged checkpoint delay to turn off the spindle checkpoint. Under normal conditions with the establishment of correct kinetochore-microtubule attachments, the checkpoint is silenced at the kinetochore due to the removal of checkpoint proteins from the kinetochore, which ultimately leads to APC/C activation [76]. For example, the normal mechanism of turning off the checkpoint after chromosomes have bioriented is through PP1’s dephosphorylation of the kinetochore protein Spc105^KNL1^ during mitosis in yeast, worms, and animal cells [3, 20, 21, 77–82]. Dephosphorylation of Spc105^KNL1^ reduces the binding affinity of Bub3, such that the kinetochore can no longer serve as a scaffold to build the MCC. Second, cells can escape the spindle checkpoint through mitotic slippage, which allows anaphase onset despite the persistence of checkpoint proteins at the kinetochore [24, 27, 29].

To distinguish between kinetochore silencing and slippage, we monitored spindle checkpoint proteins localized at the kinetochore at anaphase onset. We reasoned that with silencing, cells would disperse checkpoint proteins from the kinetochore prior to anaphase, while with slippage, cells would retain checkpoint proteins at the kinetochore during anaphase. A previous study using a similar assay found that mitotic budding yeast cells retained spindle checkpoint proteins at the kinetochore upon escaping the spindle checkpoint after the addition of microtubule depolymerizing drugs [27]. However, whether the same mechanism is used in cells that signal the checkpoint through a lack of tension or during meiosis I is not known. We tagged the spindle checkpoint protein Bub3 with three copies of mCherry (Bub3-3mCherry) in cells containing the separase biosensor (Fig 4A-B). Using live-cell imaging, we monitored Bub3-3mCherry at the kinetochore and categorized its localization at the onset of anaphase.

**Figure 4.**
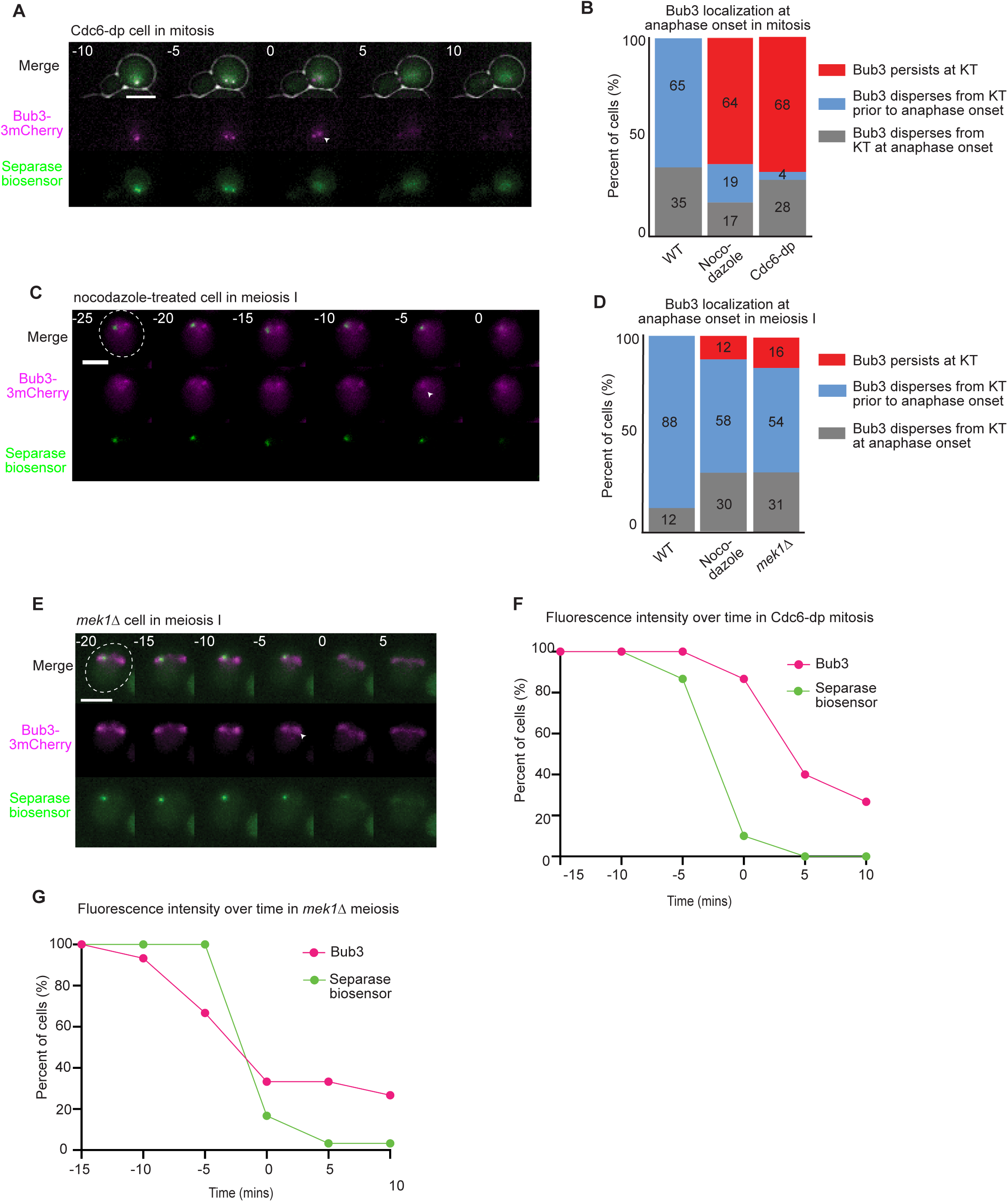
The spindle checkpoint is inappropriately silenced in meiosis I but not in mitosis. A) Representative time-lapse images show cells entering anaphase, as indicated by the dispersal of separase biosensor focus. Time 0 indicates anaphase onset. Scale bars= 5 µm. Arrowhead shows Bub3-3mcherry focus after anaphase onset. B) Quantification of Bub3-3mCherry localization at anaphase onset in mitosis. C) Representative time-lapse images of a nocodazole-treated cell. Arrowhead shows the absence of Bub3 focus at the kinetochore. Scale bars= 5 µm. D) Quantification of Bub3-3mCherry localization at anaphase onset in meiosis I. E) Representative time-lapse images of a *mek1Δ* cell entering anaphase I. Time 0 indicates anaphase onset. Arrowhead shows the absence of Bub3 focus at the kinetochore. F-G) Quantification of fluorescence intensity at the kinetochore. At each time point indicated, the fluorescence intensity of the separase biosensor and Bub3-3mcherry were measured and compared to the measurements taken at −15 minutes. Time 0 indicates anaphase onset. Graphs display the percent of cells that, at each time point, retained at least 50% of the fluorescence intensity measured 15 minutes prior to anaphase onset.

We monitored the localization of Bub3-3mCherry during mitosis, using nocodazole to disrupt kinetochore-microtubule attachments, and depletion of Cdc6 to disrupt kinetochore tension. We observed three phenotypes. First, Bub3-3mCherry dispersed from the kinetochore prior to anaphase onset. Second, Bub3-3mCherry dispersed in the same five-minute time interval in which anaphase onset occurred. Third, Bub3-3mCherry stayed localized during anaphase onset, not dispersing until after anaphase onset. When cells were treated with nocodazole in mitosis, 64% retained Bub3-3mCherry at anaphase onset, as scored by separase biosensor cleavage (Figure 4B). These results supported previous findings of mitotic slippage in the presence of nocodazole [27]. Similarly, we observed that 68% of cells depleted for Cdc6 retained Bub3-3mCherry at the kinetochore at anaphase onset, suggesting that most mitotic cells underwent mitotic slippage after prolonged checkpoint activation (Figure 4B).

We performed similar experiments in meiosis, using either nocodazole to disrupt kinetochore-microtubule attachments or deletion of *MEK1* to decrease kinetochore tension. Strikingly, with nocodazole treatment, 58% of cells dispersed Bub3-3mCherry from the kinetochore prior to anaphase I onset (Figure 4C-D). In *mek1Δ* cells, Bub3-3mCherry localized to kinetochore foci, but also to the spindle, especially at anaphase onset (Figure 4D-E). Similar to the nocodazole treated cells, the focus of Bub3 at the kinetochore dispersed from the kinetochore prior to anaphase I onset in 54% of *mek1Δ* cells. Although, we note that there is still a population on the microtubules.

To further analyze our results, we measured the fluorescence intensity of both kinetochore-localized Bub3-3mcherry and the separase biosensor foci every five minutes, from 15 minutes prior to anaphase until 10 minutes after anaphase. We plotted the percentage of cells that retained at least half of the fluorescence measured at 15 minutes prior to anaphase. In Cdc6-dp mitotic cells, 87% of cells retained at least half of their localized Bub3 fluorescence at anaphase onset (Figure 4F). In contrast, only 38% of *mek1Δ* cells retained at least half of their Bub3 fluorescence intensity prior to, or at, anaphase I onset (Figure 4G). These results are consistent with the conclusion that cells in mitosis undergo mitotic slippage, while cells in meiosis I silence the checkpoint at the kinetochore after a prolonged delay.

Because our findings differed in meiosis I and mitosis, we performed additional analyses to further scrutinize the phenotype we observed with the meiotic spindle checkpoint. We monitored Bub3-eGFP kinetochore localization in cells expressing mCherry-Tub1, scoring anaphase I onset as the time at which spindle elongation occurred in wildtype and *mek1Δ* cells (Figure S4A). Similar to wildtype, most *mek1Δ* cells dispersed Bub3-eGFP from the kinetochore prior to anaphase I onset (Figure S4B). Finally, we monitored Mad2-3GFP, another kinetochore-localized spindle checkpoint protein, in wildtype and *mek1Δ* cells. Although these cells displayed additional GFP foci, we monitored the foci that were near the SPBs and found that Mad2-3GFP dispersed from the kinetochore prior to anaphase I onset in 67% of *mek1Δ* cells (Figure S4C-E). These results suggest that unlike mitotic cells, most meiosis I cells underwent inappropriate spindle checkpoint silencing after a delay.

We next wondered if the checkpoint in meiosis II more closely resembled that of meiosis I or mitosis. To disrupt kinetochore tension in meiosis II, we deleted *SPO12*. Previous experiments showed that *spo12Δ* cells undergo meiosis I normally but fail to duplicate SPBs in meiosis II, producing two half-spindles or one weak spindle in between the two old SPBs [83–85]. We then monitored the cleavage of the centromeric Rec8-GFP at anaphase II. Similar to previous findings, the *spo12Δ* cells were delayed in cohesin cleavage, with an average duration from cohesin cleavage in anaphase I to cohesin cleavage in anaphase II of 156 ± 54 minutes (average ± SD), which is 93 minutes longer than that observed in wildtype cells (Figure S5A-C) [85]. The delay was dependent on the spindle checkpoint because disruption of *MAD3* resulted in a decreased time to anaphase II onset. We conclude that the spindle checkpoint is also less robust in meiosis II compared to mitosis.

One hypothesis for a faster anaphase II onset in meiosis II compared to mitosis is that precocious APC/C activation occurs through the meiosis-specific co-activator the APC/C, Ama1, which is active at the end of meiosis II [85]. Ama1 is impervious to the spindle checkpoint, which inhibits the APC/C coactivator Cdc20. To test this hypothesis, we deleted *AMA1* in *spo12Δ* cells and measured the duration of anaphase II onset. The *spo12Δ ama1Δ* cells had a delay with a duration similar to that of *spo12Δ* cells. Therefore, the shorter duration of the spindle checkpoint delay in meiosis II compared to mitosis was not dependent on Ama1.

To determine if the meiosis II cells underwent mitotic slippage or checkpoint silencing, we monitored Bub3-3mcherry kinetochore localization at anaphase II onset. Surprisingly, we found that 87% of cells retained Bub3-3mcherry at the kinetochore at anaphase II onset, suggesting that they underwent mitotic slippage (S4D-E). Similarly, 83% of *spo12Δ ama1Δ* cells retained Bub3-3mcherry at the kinetochore at anaphase II onset. Therefore, in meiosis II, slippage was not dependent on the activation of APC/C^Ama1^. Overall, we conclude that meiosis I has a mechanism for spindle checkpoint silencing after a prolonged delay, whereas cells in mitosis or meiosis II undergo mitotic slippage.

### Kinetochore-localized PP1 prematurely silences the spindle checkpoint in meiosis I

We hypothesized that with prolonged spindle checkpoint activity in meiosis, cells prematurely triggered the normal kinetochore checkpoint silencing pathway through PP1. The current model for spindle checkpoint silencing at the kinetochore in budding yeast is that once proper kinetochore-microtubule interactions are established, PP1 binds Spc105^Knl1^ and dephosphorylates Mps1 and Ipl^Aurora^ ^B^ substrates, releasing Bub3-Bub1 from the kinetochore (Figure 5A) [1]. When chromosomes are not properly attached, Ipl1^Aurora^ ^B^ counteracts PP1’s kinetochore binding by phosphorylating the RVSF motif on Spc105. Previous reports have shown that in vegetative cells, the s*pc105^RASA^* mutation, which disrupts PP1’s kinetochore binding, is lethal, hypothesized to be due to persistent spindle checkpoint activity [21].

**Figure 5.**
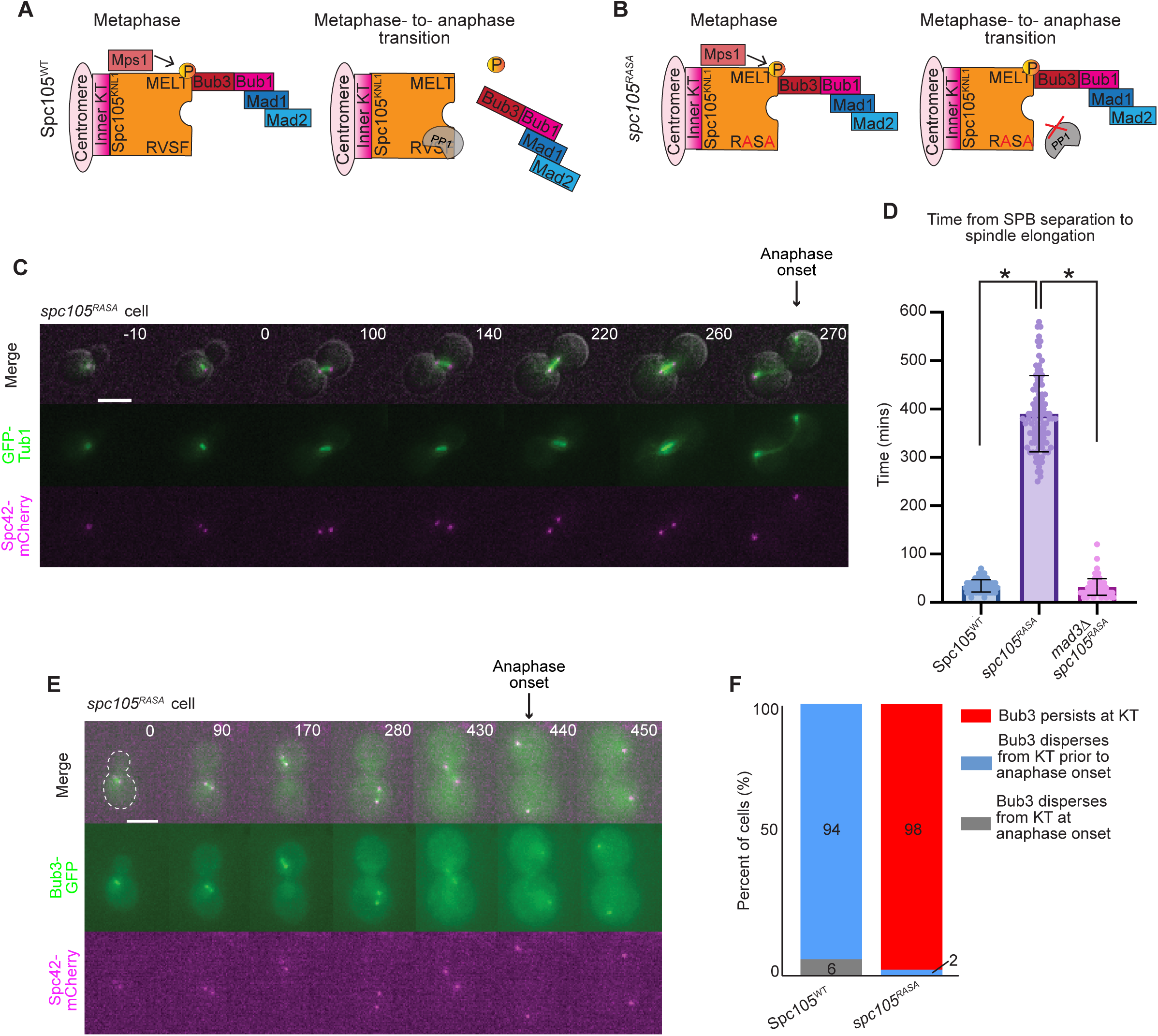
Mitotic slippage occurs in *spc105^RASA^* mitosis. A) Cartoon representation of Spc105^WT^ in metaphase (left) and at the metaphase-to-anaphase transition (right). To signal the spindle checkpoint, Mps1 phosphorylates Spc105 for binding of spindle checkpoint proteins at the kinetochore. At the metaphase-to-anaphase transition, PP1 binds Spc105 at the RVSF motif to dephosphorylate Spc105 and silence the spindle checkpoint. B) Cartoon representation of *spc105^RASA^* mutation, which prevents PP1 from binding at the RVSF motif on Spc105 and prevents PP1 from silencing the spindle checkpoint. C) Representative time-lapse images of a *spc105^RASA^*cell undergoing delayed spindle elongation. Time 0 is the time at which SPBs separate. Scale bar = 5 µm. D) Graph depicts mean duration of mitosis, measured from SPB separation to spindle elongation. n≥100 cells per genotype. Error bars show SD, * indicates statistical significance, p<0.05, Mann-Whitney test. E) Representative time-lapse images of a *spc105^RASA^*cell undergoing mitotic slippage. Time 0 is the time at which SPBs separated. Scale bar = 5 µm. F) Percentage of cells that retained Bub3-eGFP at the kinetochore during anaphase I (red), dispersed Bub3-eGFP prior to anaphase I onset (blue), and dispersed Bub3-eGFP from the kinetochore at the same time frame as anaphase I onset (gray). n≥100 cells per genotype.

We first asked if the lethality of the s*pc105^RASA^*cells in mitosis was caused by a persistent checkpoint arrest, or if the cells were able to undergo mitotic slippage. We used the anchor away system to deplete Spc105 from the nucleus in cells expressing s*pc105^RASA^* under the *SPC105* promoter. We tagged endogenous *SPC105* at its C-terminus with a FRB tag in a strain with *RPL13A* tagged with FKBP12. Addition of rapamycin allows the stable interaction between FRB and FKBP12, depleting Spc105-FRB from the nucleus but leaving Spc105^RASA^ in the nucleus. As a control, we integrated a wildtype copy of *SPC105* into the genome of strains in which we anchored away endogenous Spc105. We found that *spc105^RASA^*cells underwent anaphase onset, after a prolonged metaphase delay, with anaphase onset approximately 356 minutes later than Spc105-FRB cells expressing wildtype *SPC105* or *mad3Δ* cells expressing s*pc105^RASA^*(Figure 5C-D). The *spc105^RASA^*cells died after the first or second cell cycle following rapamycin addition. Furthermore, we observed that Bub3 remained localized at the kinetochore at anaphase onset in 98% of cells, suggesting persistent checkpoint signaling (Fig 5E-F). Therefore, we conclude that the cells that cannot localize PP1 underwent mitotic slippage after a prolonged checkpoint arrest.

To determine if PP1 is needed for the premature checkpoint silencing seen in meiosis I, we integrated mutant *spc105^RASA^* (expressed under the meiosis-specific *REC8* promoter) into a strain expressing *SPC105-FRB*. As a control, we integrated a wildtype copy of *REC8* promoted *SPC105* into the genome in strains that expressed *SPC105-FRB* (*SPC105^WT^*). The control cells with *SPC105^WT^* completed both meiotic divisions with normal timings (54 ± 20 minutes for meiosis I and 42 ± 12 minutes for meiosis II in the *SPC105^WT^* strain, compared to 45 ± 15 minutes and 51 ± 17 minutes, respectively, in a wildtype strain; Figure 6A-B).

**Figure 6.**
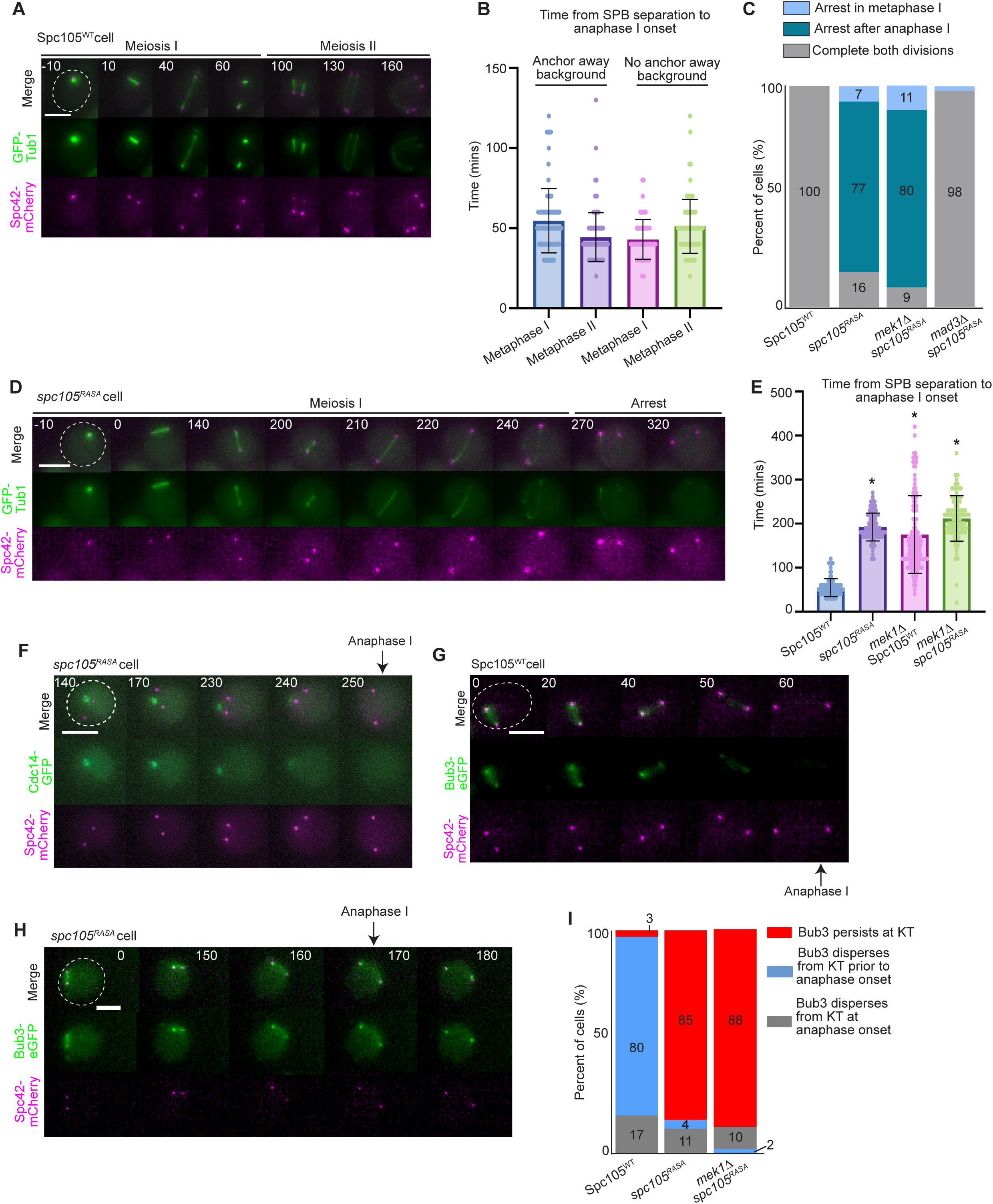
Kinetochore-localized PP1 prematurely silences the spindle checkpoint in meiosis. A-B). Time 0 indicates SPB separation. Scale bar = 5 µm. A) Representative time-lapse images of a *SPC105^WT^* cell completing both meiotic divisions. B) Graph showing the duration of metaphase I and metaphase II in *SPC105^WT^*cells with and without the anchor away background. Bars show SD. C) Phenotypes of cells that entered prometaphase I were scored. n≥100 cells per genotype. D) Representative time-lapse images of a *spc105^RASA^*cell arresting after anaphase I. E) Graph showing the mean time from SPB separation to anaphase I spindle elongation. Error bars show SD. * indicates statistical significance between wildtype and the indicated genotype as determined by Mann-Whitney test (p <0.05). F) Time-lapse images of a *spc105^RASA^* cell releasing Cdc14-GFP from nucleolus at the time of spindle elongation. Time 0 is the time at which SPBs separate. Scale bar= 5 µm. G-H) Representative time-lapse images of *SPC105^WT^* (G) and *spc105^RASA^*(H) cells. I) Percentage of cells that retained Bub3-eGFP at the kinetochore during anaphase I (red), dispersed Bub3-eGFP prior to anaphase I onset (blue), and dispersed Bub3-eGFP from the kinetochore at the same time frame as anaphase I onset (gray). n≥100 cells per genotype.

The *spc105^RASA^*cells that entered metaphase I exhibited three phenotypes: 7% arrested in metaphase I for the duration of the imaging; 77% completed anaphase I after a delay but did not progress into metaphase II; and 16% completed both meiotic divisions (Figure 6C-D). Those that progressed into anaphase I had a metaphase I duration of 192 ± 32 minutes (mean ± SD; Figure 6E). The failure to complete meiosis was due to spindle checkpoint activity; deletion of the spindle checkpoint protein *MAD3* in the *spc105^RASA^* cells resulted in 98% of cells that completed both meiotic divisions. We find that 100% of the cells that underwent anaphase I also released the phosphatase Cdc14 from the nucleolus, serving as another marker for anaphase I onset (Figure 6F). These results suggest that the cells are undergoing a transition from metaphase I to anaphase I, not just inappropriate spindle elongation.

Similar to *spc105^RASA^* cells, the *mek1Δ spc105^RASA^* cells that progressed into metaphase I also displayed the three phenotypes with 11% arrested in metaphase I for the duration of the movie, 88% arrested after anaphase I, and 9% completed both divisions (Figure 6C). Interestingly, the *mek1Δ spc105^RASA^* cells that underwent anaphase I did so after a delay that was similar in time to the *mek1Δ* delay, in that metaphase I was 212 ± 52 minutes, which is 157 minutes longer than the cells expressing wildtype *SPC105* (mean ± SD; Figure 6E, 1H). These results demonstrate that spindle checkpoint activity causes a delay but does not completely block anaphase I onset in cells that cannot localize PP1 to the kinetochore.

We were surprised to find that the *spc105^RASA^*and *mek1Δ spc105^RASA^*cells were able to undergo anaphase I after the delay. We questioned whether these cells were silencing or undergoing slippage in response to spindle checkpoint activity. We monitored the localization of Bub3-eGFP at the onset of spindle elongation. We found that 80% of cells with wildtype *SPC105* dispersed Bub3-eGFP from the kinetochore prior to anaphase I spindle elongation, as expected (Figure 6G,I). In contrast, in greater than 85% of *spc105^RASA^* and *mek1Δ spc105^RASA^* cells, Bub3 persisted at the kinetochore after spindle elongation (Figure 6H-I). These results are consistent with mitotic slippage, in that in the absence of PP1 kinetochore localization, the spindle checkpoint proteins were not released from the kinetochore, but the cells were able to undergo anaphase through another mechanism. These results suggest that in meiosis I there are two mechanisms to ensure that cells can escape a permanent checkpoint arrest: i) after a delay, cells will silence the spindle checkpoint through PP1-dependent removal of the checkpoint proteins at the kinetochore, and ii) if silencing is abrogated, cells will undergo slippage in response to persistent spindle checkpoint activity. However, most of the spc*105^RASA^* and *mek1Δ* s*pc105^RASA^* cells that escape spindle checkpoint activity were unable to enter meiosis II, suggesting that the cell cycle is mis-regulated due to persistent checkpoint activation. We conclude that with prolonged spindle checkpoint activity in meiosis I, cells use a PP1-dependent mechanism to prematurely silence the checkpoint, whereas in mitosis, cells undergo mitotic slippage.

### PP1 kinetochore binding sets the duration of the spindle checkpoint delay

When chromosomes are not properly attached to microtubules, Ipl1^Aurora^ ^B^ counteracts PP1’s kinetochore binding by phosphorylating the RVSF motif in Spc105. The phosphorylation can be abrogated by changing the RVSF motif to RVAF [21, 81]. The s*pc105^RVAF^* cells have an accelerated metaphase I and metaphase II, but not an accelerated mitosis [21, 75]. We hypothesized that if PP1 binding at the kinetochore sets the timing of checkpoint silencing, then *mek1Δ* cells expressing *spc105^RVAF^* should silence the checkpoint more quickly than *mek1Δ SPC105^WT^* cells. Indeed, we found that *mek1Δ spc105^RVAF^* cells progressed from prometaphase I to anaphase I at an average time of 106 ± 24 minutes, compared to 190 minutes in *mek1Δ SPC105^WT^* cells (Fig 7A). Furthermore, Bub3-eGFP dispersed from kinetochores before anaphase onset in 86% of *mek1Δ spc105^RVAF^* cells analyzed (Fig S4B). Thus, premature PP1 binding accelerates the time of anaphase onset through checkpoint silencing in *mek1Δ* cells. We conclude that PP1 kinetochore localization sets the duration of the spindle checkpoint arrest in meiosis.

**Figure 7.**
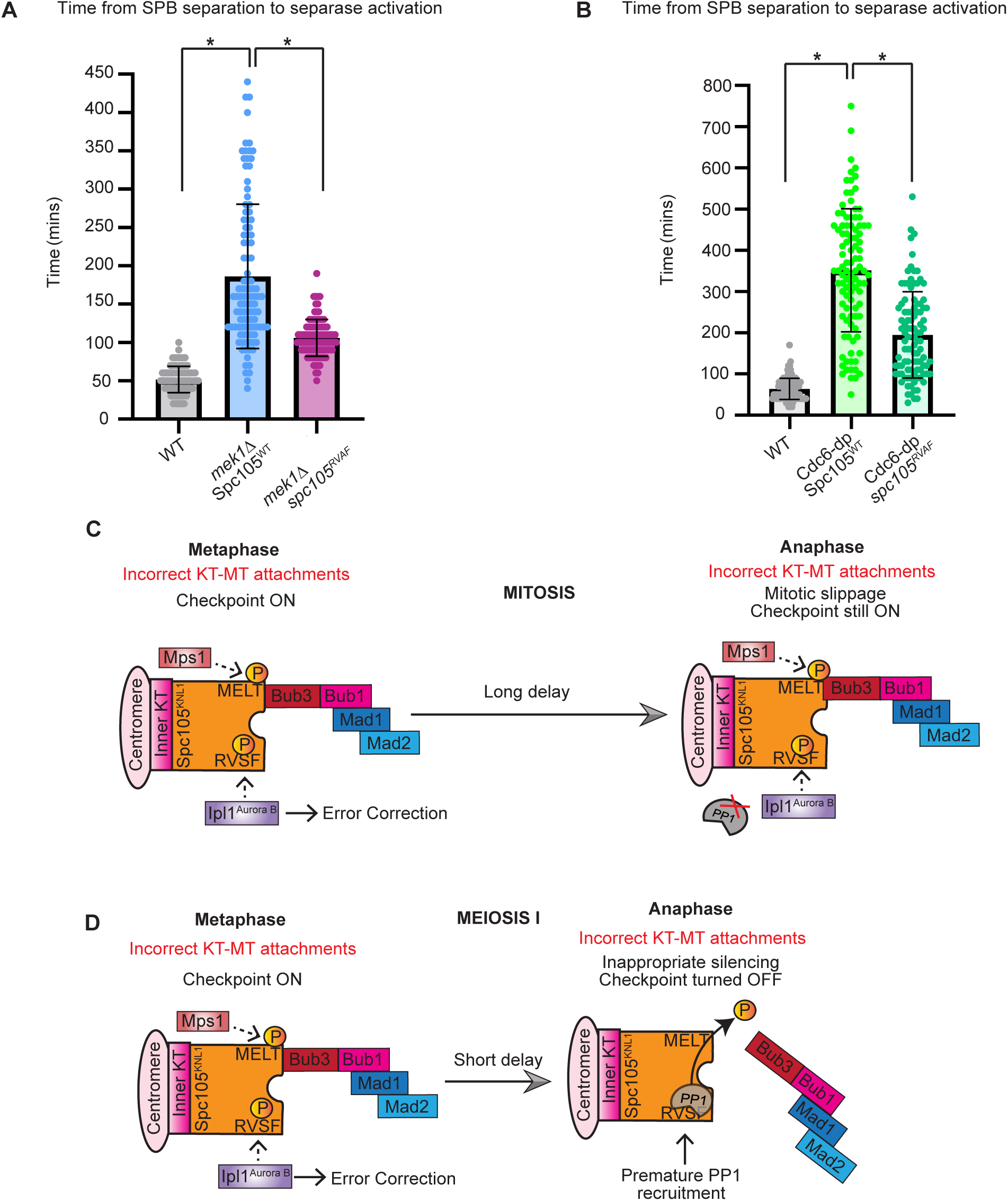
PP1 kinetochore binding sets the duration of the spindle checkpoint delay. A) Graph showing the mean time from SPB separation to separase activation in meiosis in the indicated genotypes. Error bars show SD. Asterisk indicates statistical significance relative to wildtype (p< 0.05, Mann-Whitney test). n≥100 cells per genotype. B) Graph showing the mean time from SPB separation to separase activation in mitosis in the indicated genotypes. Error bars show SD. Asterisk indicates statistical significance relative to wildtype (p< 0.05, Mann-Whitney test). n≥100 cells per genotype C-D) Model of PP1 prematurely silencing the spindle checkpoint in meiosis I, but not in mitosis. C) Activation of the spindle checkpoint in mitosis causes a relatively long delay, which cells eventually escape through mitotic slippage. D) Activation of the spindle checkpoint in meiosis causes a relatively short delay, which cells escape through a PP1-dependent checkpoint silencing. Failure of PP1 to localize at Spc105’s RVSF motif causes meiotic cells to undergo slippage. Premature PP1 kinetochore recruitment accelerates meiotic checkpoint silencing.

Although we observed that Cdc6-dp cells underwent mitotic slippage in response to a prolonged checkpoint delay, we wondered if forcing PP1’s premature kinetochore localization could accelerate the time at which Cdc6-dp cells escaped the checkpoint. We expressed *spc105^RVAF^*in Cdc6-dp cells and measured the duration of SPB separation to separase activation. We found that Cdc6-dp *spc105^RVAF^* cells were in metaphase for 195 ± 104 minutes, which is 155 minutes faster than Cdc6-dp cells (average ± SD; Figure 7B). Thus, allowing PP1 to prematurely bind the kinetochore in mitosis or meiosis reduces the duration of the spindle checkpoint delay.

## DISCUSSION

Overall, our comparison of the spindle checkpoint activity between meiotic and mitotic cells revealed interesting differences between the two types of cell division cycles. First, cells in meiosis I and meiosis II escaped spindle checkpoint activity approximately 150 minutes faster than the cells in mitosis (Figures 1, S1). These results suggest that the meiotic spindle checkpoint is less robust than the mitotic spindle checkpoint. Second, we find that cells escaped the spindle checkpoint arrest using two different mechanisms in meiosis I compared to mitosis and meiosis II. Cells in mitosis and meiosis II primarily underwent slippage to escape spindle checkpoint activity. Cells in meiosis I prematurely silenced spindle checkpoint activity through a PP1-dependent mechanism. The spindle checkpoint components were removed from the kinetochore and the signal was turned off for anaphase I onset (Figure 4B,D,F). If the kinetochore localization site of PP1was disrupted, cells instead underwent slippage (Figure 6G-H). The finding that cells in meiosis I use checkpoint silencing to escape the meiosis I arrest is interesting because normally silencing only occurs once bioriented kinetochore-microtubule attachments have been made. However, our experiments followed mutants that could not biorient kinetochores in meiosis I due to a failure to make crossovers or linkages with the homolog. These cells underwent a period of error-correction events in which the microtubules were released from kinetochores and then reattached (Figure 3A-C). After attempting to correct attachments, the cells stabilized kinetochore-microtubule attachments and turned off the checkpoint signal at the kinetochore (Figure 4B,D,F). Surprisingly, when cells expressed a mutant form of *SPC105* that prevented PP1 binding at the RVSF motif, they underwent slippage after a similar duration of spindle checkpoint delay as silencing. These results suggest that cells in meiosis I can use either mechanism to escape a prolonged checkpoint arrest in a timely manner.

PP1 is known to counteract the activities of Mps1 and Ipl^AuroraB^ [3, 20, 21, 77–82, 86–88]. We show here that both kinases are required for the full spindle checkpoint delay in *mek1Δ* cells (Figure S2). We confirm that Ipl1^AuroraB^’s kinetochore localization and error correction activity are not dampened during the meiosis I checkpoint delay (Figure 3A-C, S3). If Mps1 activity were compromised during the meiosis I checkpoint delay, then we would have expected *spc105^RASA^* mutants that cannot bind PP1 to lose Bub3 kinetochore localization during the delay. However, because we observed that Bub3-eGFP persisted at the kinetochore in *spc105^RASA^*cells, we reason that Mps1 maintained checkpoint signal throughout the delay (Figure 6H-I).

Removing Ipl1’s inhibitory phosphorylation on the RVSF residue within Spc105 (*spc105^RVAF^*), which is thought to accelerate PP1 binding, also accelerated checkpoint silencing in *mek1Δ* meiosis and Cdc6-dp mitosis (Figure 7A-B). Interestingly, when mitotic cells that do not have a persistent spindle checkpoint express *spc105^RVAF^,* there is no accelerated anaphase onset [21, 75]. However, our results show that premature PP1 binding accelerates anaphase onset when the spindle checkpoint is activated in mitosis. Because *spc105^RVAF^* mutants show an accelerated rate of anaphase onset in Cdc6-dp mitosis and *mek1Δ* meiosis, we conclude that PP1 binding to Spc105 sets the duration of checkpoint silencing (Figure 7C-D). We speculate that PP1 kinetochore localization is differentially regulated in mitosis and meiosis I to allow it to bind Spc105 earlier in meiosis I compared to mitosis (Figure 7C-D). One hypothesis is that the RVSF motif is dephosphorylated prematurely in meiosis, allowing for PP1 to bind Spc105 and silence the checkpoint.

In some organisms, the spindle checkpoint is thought to be weaker, or easier to escape, in meiosis compared to mitosis. For example, a few misaligned chromosomes in mouse oocytes do not prevent anaphase I onset, which supports the idea that the checkpoint is compromised in female meiosis [56, 59–63, 89]. However, the differences in spindle checkpoint activity are often attributed to developmental differences in the germ cells. For example, the large size of the mammalian oocyte contributes to a decrease in spindle checkpoint strength, in that the larger mouse oocytes progress through meiosis faster than smaller oocytes [56–63]. Reducing the volume of the oocyte also enhanced spindle checkpoint activity, causing a longer delay [57, 58].

Whether there are inherent differences in the duration of spindle checkpoint activity with direct comparisons between meiosis and mitosis in cells of similar sizes had not been tested. We addressed this question using budding yeast, and found that the spindle checkpoint is less robust in meiosis than in mitosis. We propose that there is a developmentally regulated mechanism to escape spindle checkpoint activity in meiosis to ensure the production of gametes. The ability to prevent a permanent checkpoint arrest is likely extremely important because cells have evolved two mechanisms in budding yeast to ensure they escape the checkpoint in meiosis I: silencing and slippage. Interestingly, in mammalian females there is some evidence that cells have evolved mechanisms to counteract a weakened meiotic spindle checkpoint and safeguard their gametes. For example, mouse oocytes produce an excess of free cyclin B1. Although cyclin B1 degradation begins early in prometaphase I, possibly due to a weakened spindle checkpoint, Cdk1-bound cyclin B1 is not targeted until anaphase onset [89]. Such overproduction of an APC/C substrate is perhaps a way by which mammals have evolved to counteract a weak meiotic spindle checkpoint. An alternative hypothesis is that the weakened spindle checkpoint is a response to the excess cyclin B1 levels. By degrading excess cyclin B1 during prolonged metaphase I, oocytes can ensure a switch-like anaphase onset, only needing CDK-bound cyclin B1 to be degraded at anaphase. In this model, it is advantageous for oocytes to have a weakened checkpoint to avoid delays beyond the developmentally programmed extended metaphase I.

Although escaping spindle checkpoint activity could lead to an increased chance of aneuploidy, evolutionarily, it may be more advantageous to the organism to produce gametes, even if there is a low probability that they are the correct ploidy, than to commit to a permanent checkpoint arrest. This mechanism would allow some viable gametes to be made rather than no viable gametes. Therefore, this work raises the exciting possibility that two mechanisms of escaping spindle checkpoint activity have evolved to ensure that cells complete meiosis, despite a lack of tension-producing kinetochore-microtubule attachments.

## MATERIALS AND METHODS

### Strains and Manipulations

*S. cerevisiae* strains used in this study are derivatives of W303 (Table S1). Gene deletions and gene tagging were performed using standard PCR-based lithium acetate transformation [90, 91]. Genotypes of transformed strains were verified by PCR. Plasmids containing the separase biosensor (P_CUP1_-LacI-Rec8-GFP:His3 and P_CUP1_-LacI-Scc1-GFP:His3) were gifts from David Morgan [41]. Strains containing P_GAL1_-*CDC6* and cdc6::ura3 were gifts from Andrew Murray [50]. Strains harboring homeologous chromosome V were S288C-derived strains and were back-crossed into W303 at least six times to obtain W303-derived strains harboring the homeologs. SLCs were introduced into yeast strains by restriction digest with BamHI, which linearizes the plasmid. All strains harboring SLCs were grown in media lacking leucine to maintain selection for SLCs. Spc105 point mutations were made by site-directed mutagenesis of a yeast integrating plasmid containing Spc105 with its endogenous promoter and terminator (∼700 bp upstream and downstream of the *SPC105* ORF) for mitosis experiments. For meiosis experiments, the same site-directed mutagenesis was performed on yeast integrating plasmids containing *SPC105* with its endogenous terminator but containing the *REC8* promoter (900 bp upstream of the *REC8* start codon). Mutant and wildtype control alleles of *SPC105* were integrated at the *LEU2* or *TRP1* locus in anchor away strains in which endogenous Spc105 was tagged with FRB at its C-terminus such that endogenous Spc105 was depleted from the nucleus upon addition of rapamycin. The anchor away strains also have *tor1-1* and *fpr1Δ* to prevent rapamycin toxicity.

### Growth conditions

All cultures were kept on a roller drum at the indicated temperature. All liquid media was supplemented with 1% tryptophan and 0.5% adenine. For all meiosis experiments, unless otherwise noted, cells were grown in 2 mL liquid media containing 1% yeast extract, 2% peptone, and 2% dextrose (YPD) for 10-24 hours at 30° C, then 40 μL of this culture was transferred to 2 mL of 1% yeast extract, 2% peptone, and 2% potassium acetate (YPA) for 12-15 hours at 30°C. Cells were washed twice with water and transferred to 1% potassium acetate (1% KAc) for incubation at 25° C. Time-lapse imaging of meiosis, except *NDT80*-in strains, began at 6-8 hours post-KAc transfer. For Spc105 anchor away movies, rapamycin was added to a final concentration of 1μM 6-7 hours after KAc transfer such that Spc105 remained unperturbed for prophase I. *P_GAL1,10_-NDT80 GAL4-ER* strains were kept in 1% KAc at 25° C for 10-12 hours, before release from prophase I by addition of beta-estradiol to a final concentration of 1mM. Cells harboring a SLC were grown in media lacking leucine to maintain selection for the SLC. These cells were prepared for imaging by inoculating 2 mL of 2XSC+glucose –leucine (0.67% yeast nitrogen base without amino acids, 0.2% dropout mix containing all amino acids except leucine, 2% glucose) and letting culture grow for 10-24 hours at 30° C. 40 μL of this culture was transferred to 2 mL 2XSCA - leucine (0.67% yeast nitrogen base without amino acids, 0.2% dropout mix containing all amino acids except leucine, 2% potassium acetate) for 12-16 hours at 30° C, then washed twice with water before resuspending in 1% KAc for 6-8 hours of incubation at 25° C.For all mitosis experiments, unless otherwise noted, cells were grown overnight in synthetic complete media (2XSC + glucose). For nocodazole experiments, cells were grown in 2XSC + 1% peptone, because the presence of peptone increases the effect of nocodazole in synthetic media [27, 92]. For all Cdc6-dp experiments, cells were grown in 2XSC+ 20% galactose until time of experiment.

### Time-lapse microscopy

Time-lapse imaging was performed as follows, unless otherwise noted. 200 μL of cells were concentrated and spread on a 35 mm x 40 mm coverslip coated with 1 mg/mL Concanavilin A (ConA) and fitted to a chamber. A 5% agar plug was made by cutting off the tip of an Eppendorf tube and pipetting ∼100 μL of melted 5% agar into the tube sitting on a glass slide. The dried agar plug was placed on top of the cells sitting on the ConA for 12 mins, such that the cells had time to adhere to the ConA. The rest of the Kac (or 2XSC for mitotic cells) culture (∼ 2 mL) was spun down, and the supernatant was added drop-wise to the chamber in order to gently float and remove the agar pad. After cells were adhered to the coverslip and the chamber filled with medium, the chamber was immediately placed on the microscope for imaging. Two microscopes were used for this study. Images were acquired on a Delta Vision Elite (eDV) microscope, equipped with a PCO Edge5.5 sCMOS camera and Olympus 60X oil-immersion objective lens (Plan Apo N, 1.42). Cells were imaged at room temperature using FITC and mCherry filters. FITC exposure was at 2% for 15-25 msec and mCherry exposure was at 2-5% for 100 msec. Brightfield images were acquired at 5% intensity for 50 msec. Images were acquired using the SoftWorx Version 7.0 software (G.E. Healthcare), and data analysis was performed using Fiji and Image J. Nocodazole and Spc105 mitosis experiments were performed using a Nikon Ti2 microscope equipped with a Photometrics camera and 60X oil immersion objective lens. GFP and Ruby filters were used. Images were acquired with exposure times of 20-40 msec for GFP and Ruby filters with neutral density (ND) filters transmitting 2-5% of light intensity. Brightfield images were acquired at 70 msec at 5% ND. Images were acquired and data were analyzed using NIS Elements Viewer Version 4.20 software.

All images were collected with 5, 1-1.2-μM z-stacks, in 10-minute time increments, for 10-12 hours, unless otherwise noted. For experiments on the Nikon Ti2, NIS elements was used for analysis. Fiji software (NIH) was used to create final images with adjustment of brightness and contrast.

### Image Analysis and Fluorescence Intensity Measurement

Movie analysis was conducted in Fiji software. Unless otherwise noted, z-stacks were combined into a single maximum-intensity projection for analysis and time-lapse images.

Fluorescence intensity measurement of Bub3-3mcherry and the separase biosensor focus was done with Fiji. Images were projected in SUM slices, and a small circle was drawn around each focus at the time points indicated. A background circle was drawn just beside the measured point for each channel, in each cell, at each time point, and subtracted from the fluorescence intensity measurement to generate each value. Fluorescence intensity measurement of Ipl1-3GFP and Mtw1-mRuby2 was performed using SUM slice projection of images. A small circle was drawn around Mtw1-mRuby2 using the circle tool in Fiji, and the fluorescence intensity of the mCherry channel and the GFP channel within the drawn circle were recorded.

### Spindle length measurement

Spindle length was calculated by identifying the location of one SPB in a single, 1 μm z-stack out of a 5-stack image. The opposite SPB was detected in the same way, and the number of 1 μm stacks between the two SPBs was noted. A line was draw between the two SPBs and the Pythagorean theorem was used to calculate the distance between the two SPBs.

### Cell volume measurement

Cell volume was measured using BudJ: an ImageJ plugin to analyze fluorescence microscopy images of budding yeast cells (http://www.ibmb.csic.es/home/maldea; [93]). Time-lapse imaging files were cropped such that the frames from 20 mins prior to anaphase until 30 minutes after anaphase were included in the analysis (the data reported are from 10 mins prior to anaphase onset). For cells in mitosis, the volume reported is the volume of the mother plus the volume of the bud.

### Cdc6 depletion

For Cdc6-dp experiments, cells were grown overnight in 2X SC + galactose medium and diluted 1:20 the following day. After growth for 3 h, cells were spun down, rinsed 3 times with 2X SC + glucose, and resuspended in 2X SC + glucose containing 20 μM copper sulfate to induce expression of the separase biosensor. Cells were immediately spread on a coverslip containing Concanavilin A (ConA) and resuspended in the 2X SC + glucose containing 20 μM copper sulfate. For analysis, only the cells which were in S phase through early M phase at the start of imaging were analyzed, as determined by spindle length. These cells were monitored through the rest of the cell cycle, and only the following cell cycle was analyzed to allow for depletion of Cdc6 and to avoid analyzing cells which had undergone an entire cell cycle without Cdc6.

### Nocodazole treatment

For all nocodazole (Sigma M1404) experiments, 30 mg/mL stocks in DMSO were kept at −20C in 20 μL aliquots to avoid multiple freeze-thaw cycles of the drug. Because nocodazole interferes with SPB separation, the drug was not added until two SPBs were observed. Additionally, image analysis of all nocodazole-treated cells and their respective controls was performed such that the time at which the imagining began was regarded as time 0.

For mitosis experiments, cells were grown in 2X SC + 1% peptone. Addition of peptone has been shown to increase the effect of nocodazole in synthetic media [27, 92]. An overnight culture of cells was grown in 2XSC + 1% peptone+ 20 μM copper sulfate and diluted to 1:20 the following day. After 3 h of growth cells were arrested with 25 μM α-factor (Zymo Research) for 3 h. Cells were washed 5 times with 2X SC + 1% peptone + 20 µM copper sulfate to release from α -factor arrest. Following release, cells were grown for 30 mins at 30° C, and imaged under the microscope to confirm separation of SPBs. Cells were then immediately re-suspended in 2X SC + 1% peptone + 20 μM copper sulfate to which nocodazole was added drop-wise to a final concentration of 15 ug/mL. Cells were immediately spread on a coverslip containing ConA for imaging, and 2X SC + 1% peptone containing 15 μg/mL nocodazole and 20 μM copper sulfate was added to the chamber surrounding the coverslip.

For all meiosis nocodazole experiments, pre-conditioned media was prepared by taking a diploid strain through the sporulation process (12 h YPD, 12 h YPD, 24 h 1% KAc). The sporulation culture was then spun down and the supernatant filter sterilized. 1-2 h before the start of imaging, beta-estradiol was added to a final concentration of 1 mM, copper sulfate was added to a final concentration of 20µM, and nocodazole was added dropwise to a final concentration of 30 μg/mL. All meiosis nocodazole experiments were performed in cells harboring the *NDT80* gene under control of the *GAL1,10* promoter with the transcriptional activator *GAL4-ER* integrated into the genome, such that addition of beta-estradiol releases Gal4 into the nucleus to activate Ndt80 expression. Cells were grown in YPD for 8-12 h, after which 40 μL of culture was transferred to 3 mL YPA for 12-15 h, and then transferred to 3 mL 1% KAc for 12 h to induce a prophase I arrest. Cells were released from prophase I arrest by addition of beta-estradiol to a final concentration of 1 mM. Just after SPB separation was detected (∼80 mins after release from prophase I), cells were resuspended in pre-conditioned media described above (1% KAc with 1 mM beta-estradiol, 20 µM copper sulfate, and 30 μg/mL nocodazole, and immediately spread on a coverslip containing ConA for imaging. 2 mL of the pre-conditioned 1% KAc + 30 μg/mL nocodazole + 20 μM copper sulfate was added to the chamber surrounding the coverslip.

## ACKNOWLEDGEMENTS

We thank the Lacefield lab, Brian Calvi, and Claire Walczak for insightful comments on the manuscript. We thank Andrew Murray and David Morgan for strains and plasmids. We thank the Light Microscopy and Imaging Center at Indiana University, especially Jim Powers for assistance. Our research was supported by NIH grant GM105755 to SL.

## Supplemental Figure Legends

**Figure S1.**
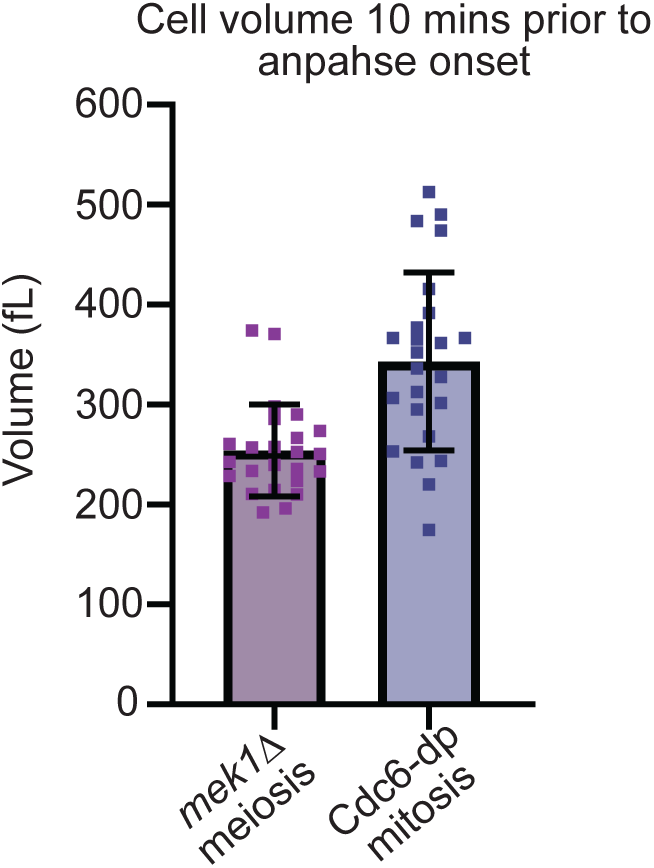
Mitotic and meiotic cells have similar volumes at anaphase onset after a spindle checkpoint delay. A) Graph showing cell volume measured 10 minutes prior to anaphase onset in *mek1Δ* meiosis and Cdc6-dp mitosis. Error bars indicate SD. n= 25 cells per genotype.

**Figure S2.**
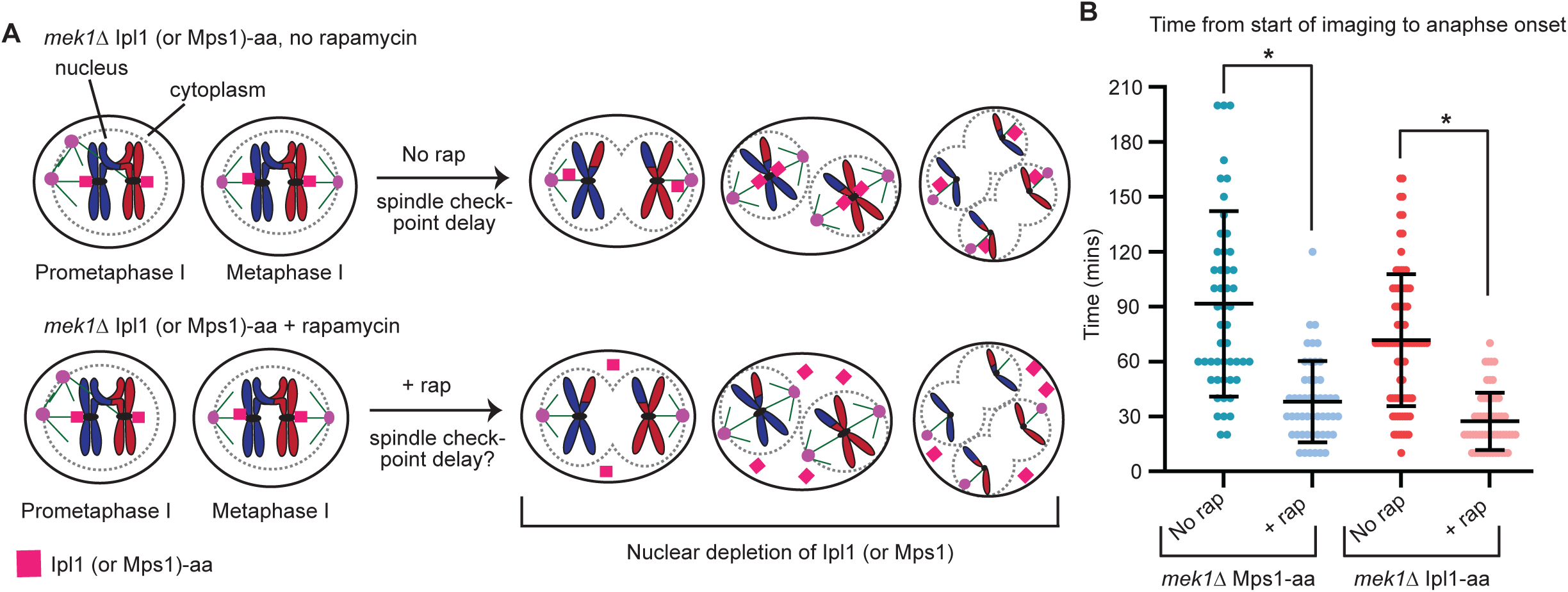
Mps1 and Ipl1 are required for the spindle checkpoint delay in *mek1Δ* meiosis. A) Cartoon of experimental design. Ipl1-aa *mek1Δ* and Mps1-aa *mek1Δ* strains progressed into metaphase I, and were then treated with rapamycin to deplete Ipl1 or Mps1 from the nucleus. Live-cell imaging began immediately following rapamycin treatment, and time from start of imaging to anaphase spindle elongation was measured. B) Graph showing time from start of imaging to anaphase onset in individual cells. Center bar is mean, and error bars represent SD. Rap = Rapamycin; aa = anchor away. n≥50 cells per genotype. Asterisk shows statistical significance between the indicated conditions (Welch’s t-test, p < 0.05).

**Figure S3.**
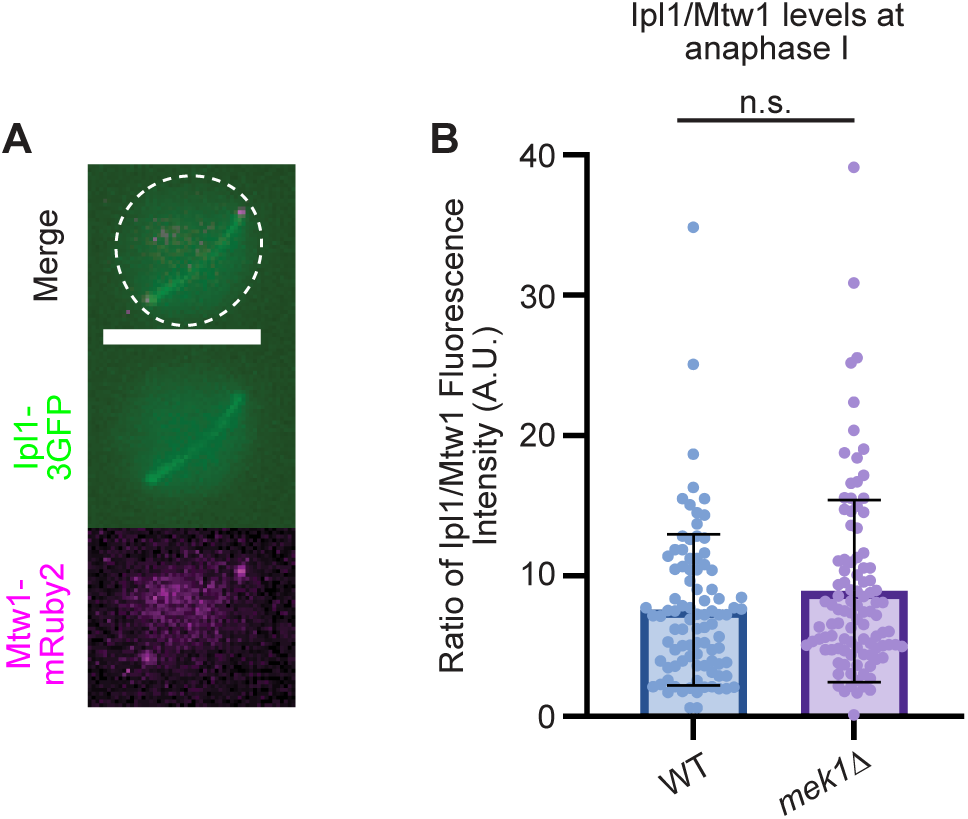
WT and *mek1Δ* cells have similar levels of Ipl1 at the kinetochore at anaphase I onset. A) Representative images of a *mek1Δ* cell at anaphase I onset. Measurements were made by drawing circles around the Mtw1-mRuby2 focus (representing the kinetochore) and recording both Mtw1-mRuby2 and Ipl1-3GFP fluorescence intensity. Scale bar = 5 μm. B) Quantification of Ipl1/Mtw1 fluorescence intensity. 90 or more cells per genotype were scored, and the mean is plotted. Mann-Whitney test was performed and showed no statistically significant difference between WT and *mek1*Δ. Error bars show SD.

**Figure S4.**
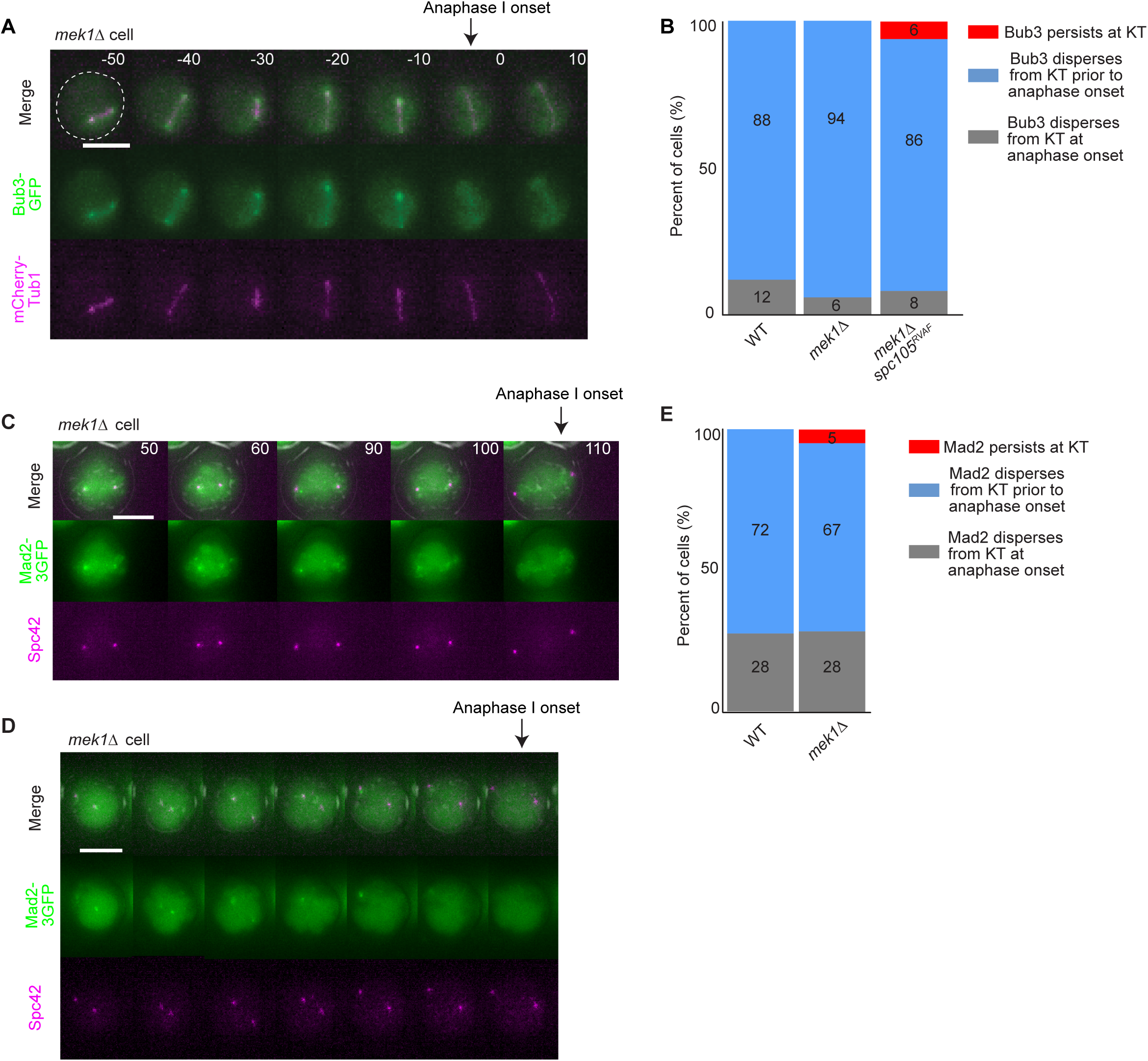
Bub3-eGFP and Mad2-3GFP disperses from the kinetochore prior to anaphase I spindle elongation. A) Time-lapse images of a *mek1Δ* cell show Bub3-eGFP and mCherry-Tub1 during meiosis I. Time 0 is set as the time frame in which anaphase onset occurs. Scale bar = 5 µm. B) Quantification of Bub3-eGFP dispersal in WT and *mek1Δ* cells. n≥100 cells per genotype. C) Time lapse images of a *mek1Δ* cell in which Mad2-3GFP disperses at anaphase I onset. Time 0 is the time at which SPB separation occurs. Scale bar = 5 µm. D) Time lapse images of a *mek1Δ* cell in which Mad2-3GFP disperses from the kinetochore prior to anaphase onset. Scale bar = 5 µm. E) Quantification of Mad2-3GFP dispersal in WT and *mek1Δ* cells. n≥100 cells per genotype.

**Figure S5.**
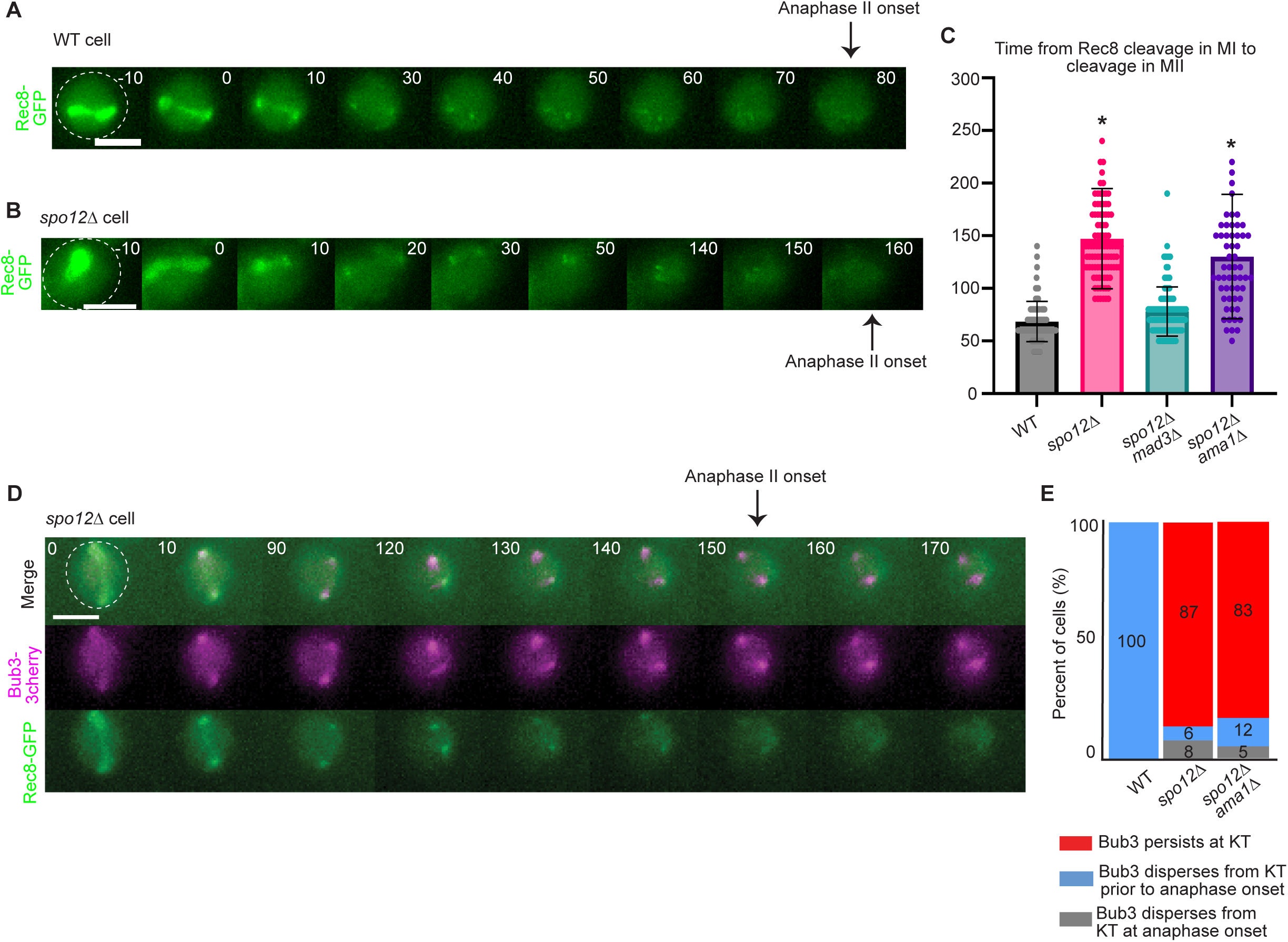
Cells lacking kinetochore tension in meiosis II undergo slippage. A-B) Representative time-lapse images of WT and *spo12Δ* cells expressing Rec8-GFP. Time 0 is the time at which Rec8 cleavage initiates in meiosis I. Scale bar= 5 µm. Error bars show SD. C) Quantification of the duration from Rec8 cleavage in meiosis I to Rec8 cleavage in meiosis II. Statistical significance was determined by Mann-Whitney test (p < 0.05). Asterisk indicates statistically significant difference between wildtype and the indicated genotypes. n≥ 100 cells per genotype. D) Time-lapse images of cells with Rec8-GFP and Bub3-3mcherry. Time 0 is the time at which Rec8 is cleaved in meiosis I. E) Quantification of Bub3-3mCherry dispersal. n≥100 cells per genotype.

**Table S1.** Budding yeast strains used in this study.

| Strain number: | Strain genotype: |
| --- | --- |
| LY3022 | MATa/$\alpha$ Spc42-mCherry:kanMX6/+, P<sub>TUB1</sub>GFP-TUB1:LEU2/ P<sub>TUB1</sub>GFP-TUB1:LEU2, ZIP1-GFP/+ |
| LY5451 | MAT a, Spc42-mCherry:HphMX, LacO:TRP1, P<sub>CUP1</sub>- GFP-Scc1-LacI:HIS3 |
| LY5512 | MAT a, Spc42-mCherry:HphMX, LacO:TRP1, P<sub>CUP1</sub>- GFP-Scc1-LacI:HIS3, mad3::KanMX |
| LY8248 | MAT a/$\alpha$, Spc42-mCherry:HphMX /+, P<sub>CUP1</sub>- GFP-Rec8-LacI:HIS3/+, LacO:TRP1 at 800 kb from CEN4/ LacO:TRP1 at 800 kb from CEN4, P<sub>GAL1-1</sub>-Ndt80:TRP1/ P<sub>GAL1-1</sub>-Ndt80:TRP1, Gal4-ER:URA3/ Gal4-ER:URA3 |
| LY8312 | MAT a/$\alpha$, Spc42-mCherry:HphMX /+, P<sub>CUP1</sub>- GFP-Rec8-LacI:HIS3/+, LacO:TRP1 at 800 kb from CEN4/ LacO:TRP1 at 800 kb from CEN4, P<sub>GAL1-1</sub>-Ndt80:TRP1/ P<sub>GAL1-1</sub>-Ndt80:TRP1, Gal4-ER:URA3/Gal4-ER:URA3, mad3::KanMX/mad3::KanMX |
| LY5657 | MAT a/$\alpha$, Spc42-mCherry:HphMX /+, LacO:TRP1/LacO:TRP1, P<sub>CUP1</sub>-GFP-Scc1-LacI:HIS3/ P<sub>CUP1</sub>-GFP-Scc1-LacI:HIS3 |
| LY10083 | MAT a/$\alpha$, Spc42-mCherry:HphMX /+, LacO:TRP1/LacO:TRP1, P<sub>CUP1</sub>-GFP-Scc1-LacI:HIS3/ P<sub>CUP1</sub>-GFP-Scc1-LacI:HIS3, mad3::KanMX/mad3::KanMX |
| LY5661 | MAT a/$\alpha$, Spc42-mCherry:HphMX /+, cdc6::HISG/cdc6::HISG, P<sub>GAL1-1</sub>-UbiCdc6:URA3/ P<sub>GAL1-1</sub>-UbiCdc6:URA3, PDS1:Pds1myc18:Leu2/pds1:Pds1myc18:Leu2, LacO:TRP1/ LacO:TRP1, P<sub>CUP1</sub>-GFP-Scc1-LacI:HIS3/ P<sub>CUP1</sub>-GFP-Scc1-LacI:HIS3 |
| LY10033 | MAT a/$\alpha$, Spc42-mCherry:HphMX /+, cdc6::HISG/cdc6::HISG, P<sub>GAL1-1</sub>-UbiCdc6:URA3/ P<sub>GAL1-1</sub>-UbiCdc6:URA3, PDS1:Pds1myc18:Leu2/pds1:Pds1myc18:Leu2, LacO:TRP1/ LacO:TRP1, P<sub>CUP1</sub>-GFP-Scc1-LacI:HIS3/ P<sub>CUP1</sub>-GFP-Scc1-LacI:HIS3, mad3::KanMX/mad3::KanMX |
| LY5658 | MAT a/$\alpha$, Spc42-mCherry:HphMX /+, P<sub>CUP1</sub>- GFP-Rec8-LacI:HIS3/ P<sub>CUP1</sub>- GFP-Rec8-LacI:HIS3, LacO:TRP1 at 800 kb from CEN4/ LacO:TRP1 at 800 kb from CEN4 |
| LY8211 | MAT a/$\alpha$, Spc42-mCherry:HphMX /+, P<sub>CUP1</sub>- GFP-Rec8-LacI:HIS3/ P<sub>CUP1</sub>- GFP-Rec8-LacI:HIS3, LacO:TRP1 at 800 kb from CEN4/ LacO:TRP1 at 800 kb from CEN4, mad3::KanMX/mad3::KanMX |
| LY5659 | MAT a/$\alpha$, Spc42-mCherry:HphMX /+, P<sub>CUP1</sub>- GFP-Rec8-LacI:HIS3/ P<sub>CUP1</sub>- GFP-Rec8-LacI:HIS3, LacO:TRP1 at 800 kb from CEN4/ LacO:TRP1 at 800 kb from CEN4, spo11::NatMX/spo11::NatMX, spo11-Y135F:URA3/ spo11-Y135F:URA3 |
| LY5660 | MATa/$\alpha$, Spc42-mCherry:HphMX /+, P<sub>CUP1</sub>- GFP-Rec8-LacI:HIS3/ P<sub>CUP1</sub>- GFP-Rec8-LacI:HIS3, LacO:TRP1 at 800 kb from CEN4/ LacO:TRP1 at 800 kb from CEN4, spo11::NatMX/spo11::NatMX, spo11-Y135F:URA3/ spo11-Y135F:URA3, mad3::KanMX/mad3::KanMX |
| LY10068 | MAT a/$\alpha$, Spc42-mCherry:HphMX /+, P<sub>CUP1</sub>- GFP-Rec8-LacI:HIS3/ P<sub>CUP1</sub>- GFP-Rec8-LacI:HIS3, LacO:TRP1 at 800 kb from CEN4/ LacO:TRP1 at 800 kb from CEN4, mek1::KanMX/mek1::KanMX |
| LY10082 | MATa/$\alpha$, Spc42-mCherry:HphMX /+, P<sub>CUP1</sub>- GFP-Rec8-LacI:HIS3/ P<sub>CUP1</sub>- GFP-Rec8-LacI:HIS3, LacO:TRP1 at 800 kb from CEN4/ LacO:TRP1 at 800 kb from CEN4, mek1::NatMX/mek1::NatMX, mad3::KanMX/mad3::KanMX |
| LY8140 | MATa/$\alpha$, Spc42-mCherry:HphMX /+, P<sub>CUP1</sub>- GFP-Rec8-LacI:HIS3/ P<sub>CUP1</sub>- GFP-Rec8-LacI:HIS3, LacO:TRP1 at 800 kb from CEN4/ LacO:TRP1 at 800 kb from CEN4, Chrom V carlsbergensis[ivl1::pRM12[NatR:LacO]]/ Chrom V cerevisiae |
| LY8264 | MATa/$\alpha$, Spc42-mCherry:HphMX /+, P<sub>CUP1</sub>- GFP-Rec8-LacI:HIS3/ P<sub>CUP1</sub>- GFP-Rec8-LacI:HIS3, LacO:TRP1 at 800 kb from CEN4/ LacO:TRP1 at 800 kb from CEN4, Chrom V carlsbergensis[ivl1::pRM12[NatR:LacO]]/ Chrom V cerevisiae, mad3::KanMX/mad3::KanMX |
| LY8210 | MATa/$\alpha$, Spc42-mCherry:HphMX /+, P<sub>CUP1</sub>- GFP-Rec8-LacI:HIS3/ P<sub>CUP1</sub>- GFP-Rec8-LacI:HIS3, LacO:TRP1 at 800 kb from CEN4/ LacO:TRP1 at 800 kb from CEN4, SLC:V141-LacO:Leu2 |
| LY8212 | MATa/$\alpha$, Spc42-mCherry:HphMX /+, P<sub>CUP1</sub>- GFP-Rec8-LacI:HIS3/ P<sub>CUP1</sub>- GFP-Rec8-LacI:HIS3, LacO:TRP1 at 800 kb from CEN4/ LacO:TRP1 at 800 kb from CEN4, mad3::KanMX/mad3::KanMX, SLC:V141-LacO:Leu2 |
| LY8156 | MATa/$\alpha$, Spc42-mCherry:HphMX /+, P<sub>CUP1</sub>- GFP-Rec8-LacI:HIS3/ P<sub>CUP1</sub>- GFP-Rec8-LacI:HIS3, LacO:TRP1 at 800 kb from CEN4/ LacO:TRP1 at 800 kb from CEN4, Chrom V carlsbergensis[ivl1::pRM12[NatR:LacO]]/ Chrom V cerevisiae, SLC:V141-LacO:Leu2 |
| LY8303 | MATa/$\alpha$, Spc42-mCherry:HphMX /+, P<sub>CUP1</sub>- GFP-Rec8-LacI:HIS3/ P<sub>CUP1</sub>- GFP-Rec8-LacI:HIS3, LacO:TRP1 at 800 kb from CEN4/ LacO:TRP1 at 800 kb from CEN4, Chrom V carlsbergensis[ivl1::pRM12[NatR:LacO]]/ Chrom V cerevisiae, mad3::KanMX, mad3::KanMX, SLC:V141-LacO:Leu2 |
| LY4415 | MATa/$\alpha$, Spc42-mCherry: HphMX/+, P<sub>CUP1</sub>-LACI-GFP:HIS3/ P<sub>CYC1</sub>-LACI-GFP:HIS3, LacO:TRP/ LacO:TRP |
| LY4568 | MATa/$\alpha$, Spc42-mCherry: HphMX/+, P<sub>CUP1</sub>-LACI-GFP:HIS3/ P<sub>CUP1</sub>-LACI-GFP:HIS3, LacO:TRP/ LacO:TRP, mad3::KanMX/mad3::KanMX |
| LY4515 | MATa/$\alpha$, Spc42-mCherry: HphMX/+, P<sub>CUP1</sub>-LACI-GFP:HIS3/ P<sub>CUP1</sub>-LACI-GFP:HIS3, LacO:TRP/ LacO:TRP, mek1::KanMX/mek1::KanMX |
| LY4607 | MATa/$\alpha$, Spc42-mCherry: HphMX/+, P<sub>CUP1</sub>-LACI-GFP:HIS3/ P<sub>CUP1</sub>-LACI-GFP:HIS3, LacO:TRP/ LacO:TRP, mek1::KanMX/mek1::KanMX, mad3::KanMX/mad3::KanMX |
| LY10086 | MATa/$\alpha$, P<sub>CUP1</sub>- GFP-Rec8-LacI:HIS3/ P<sub>CUP1</sub>- GFP-Rec8-LacI:HIS3, LacO:TRP1 at 800 kb from CEN4/ LacO:TRP1 at 800 kb from CEN4, bub3::Leu2/bub3::Leu2, trp1::Bub3-3mCherry:TRP1/ trp1::Bub3-3mCherry:TRP1 |
| LY10067 | MATa/$\alpha$, P<sub>CUP1</sub>- GFP-Rec8-LacI:HIS3/ P<sub>CUP1</sub>- GFP-Rec8-LacI:HIS3, LacO:TRP1 at 800 kb from CEN4/ LacO:TRP1 at 800 kb from CEN4, bub3::Leu2/bub3::Leu2, trp1::Bub3-3mCherry:TRP1/ trp1::Bub3-3mCherry:TRP1, mek1::KanMX/mek1::KanMX |
| LY10066 | MATa/$\alpha$, P<sub>CUP1</sub>- GFP-Scc1-LacI:HIS3/ P<sub>CUP1</sub>- GFP-Scc1-LacI:HIS3, LacO:TRP1 at 800 kb from CEN4/ LacO:TRP1 at 800 kb from CEN4, bub3::Leu2/bub3::Leu2, |
|  | trp1::Bub3-3mCherry:TRP1/ trp1::Bub3-3mCherry:TRP1 |
| LY10036 | MATa/$\alpha$, P<sub>CUP1</sub>- GFP-Scc1-LacI:HIS3/ P<sub>CUP1</sub>- GFP-Scc1-LacI:HIS3, LacO:TRP1 at 800 kb from CEN4/ LacO:TRP1 at 800 kb from CEN4, bub3::Leu2/bub3::Leu2, trp1::Bub3-3mCherry:TRP1/ trp1::Bub3-3mCherry:TRP1, cdc6::HISG/cdc6::HISG, P<sub>GAL1-1</sub>-UbiCdc6:URA3/ P<sub>GAL1-1</sub>-UbiCdc6:URA3, PDS1:Pds1myc18:Leu2/pds1:Pds1myc18:Leu2 |
| LY9241 | MATa/$\alpha$, P<sub>CUP1</sub>- GFP-Rec8-LacI:HIS3/ P<sub>CUP1</sub>- GFP-Rec8-LacI:HIS3, LacO:TRP1 at 800 kb from CEN4/ LacO:TRP1 at 800 kb from CEN4, bub3::Leu2/bub3::Leu2, trp1::Bub3-3mCherry:TRP1/ trp1::Bub3-3mCherry:TRP1, P<sub>Gal1</sub>-Ndt80:TRP1/ P<sub>Gal1</sub>-Ndt80:TRP1, Gal4-ER:URA3/ Gal4-ER:URA3 |
| LY9417 | MATa/$\alpha$, tor1-1:HIS3/tor1-1:HIS3, fpr1::natMX4/fpr1::natMX4, RPL13A-2XFKBP12:TRP1/RPL13A-2XFKBP12:TRP1, Spc42-mCherry:HphMX/+, P<sub>TUB1</sub>GFP-TUB1:URA3/P<sub>TUB1</sub>GFP-TUB1:URA3, Spc105-FRB:KanMX/Spc105-FRB:KanMX, P<sub>Spc105</sub>-Spc105:LEU2/P<sub>Spc105</sub>-Spc105:LEU2 |
| LY10085 | MATa/$\alpha$, tor1-1:HIS3/tor1-1:HIS3, fpr1::natMX4/fpr1::natMX4, RPL13A-2XFKBP12:TRP1/RPL13A-2XFKBP12:TRP1, Spc42-mCherry:HphMX/+, P<sub>TUB1</sub>GFP-TUB1:URA3/P<sub>TUB1</sub>GFP-TUB1:URA3, Spc105-FRB:KanMX/Spc105-FRB:KanMX, P<sub>Spc105</sub>-spc105<sup>RASA</sup>:LEU2/P<sub>Spc105</sub>-spc105<sup>RASA</sup>:LEU2 |
| LY10087 | MATa/$\alpha$, tor1-1:HIS3/tor1-1:HIS3, fpr1::natMX4/fpr1::natMX4, RPL13A-2XFKBP12:TRP1/RPL13A-2XFKBP12:TRP1, Spc42-mCherry:HphMX/+, P<sub>TUB1</sub>GFP-TUB1:URA3/P<sub>TUB1</sub>GFP-TUB1:URA3, Spc105-FRB:KanMX/Spc105-FRB:KanMX, P<sub>Spc105</sub>-spc105<sup>RASA</sup>:LEU2/P<sub>Spc105</sub>-spc105<sup>RASA</sup>:LEU2, mad3::KanMX/mad3::KanMX |
| LY9533 | MATa/$\alpha$, tor1-1:HIS3/tor1-1:HIS3, fpr1::natMX4/fpr1::natMX4, RPL13A-2XFKBP12::loxP /RPL13A-2XFKBP12::loxP, Spc42-mCherry:HphMX/+, Bub3-eGFP:TRP1/ Bub3-eGFP:TRP1, Spc105-FRB:KanMX/Spc105-FRB:KanMX, P<sub>Spc105</sub>-Spc105:LEU2/P<sub>Spc105</sub>-Spc105:LEU2 |
| LY10084 | MATa/$\alpha$, tor1-1:HIS3/tor1-1:HIS3, fpr1::natMX4/fpr1::natMX4, RPL13A-2XFKBP12::loxP /RPL13A-2XFKBP12::loxP, Spc42-mCherry:HphMX/+, Bub3-eGFP:TRP1/ Bub3-eGFP:TRP1, Spc105-FRB:KanMX/Spc105-FRB:KanMX, P<sub>Spc105</sub>-Spc105<sup>RASA</sup>:LEU2/P<sub>Spc105</sub>-Spc105<sup>RASA</sup>:LEU2 |
| LY3022 | MATa/$\alpha$ Spc42-mCherry:kanMX6/+, P<sub>TUB1</sub>GFP-TUB1:LEU2/ P<sub>TUB1</sub>GFP-TUB1:LEU2, ZIP1-GFP/+ |
| LY5855 | MATa/$\alpha$, tor1-1::HIS3/tor1-1, fpr1::NatMX/fpr1::natMX4, RPL13A-2XFKBP12:TRP1/RPL13A-2XFKBP12:TRP1, Spc42-mCherry:HphMX/+, P<sub>TUB1</sub>GFP-TUB1:URA3/P<sub>TUB1</sub>GFP-TUB1:URA3, Spc105-FRB:KanMX/Spc105-FRB:KanMX, P<sub>Rec8</sub>-Spc105:LEU2/P<sub>Rec8</sub>-Spc105:LEU2 |
| LY10110 | MATa/$\alpha$, tor1-1:HIS3/tor1-1:HIS3, fpr1::NatMX/fpr1::natMX4, RPL13A-2XFKBP12:TRP1/RPL13A-2XFKBP12:TRP1, Spc42-mCherry:HphMX/+, P<sub>TUB1</sub>GFP-TUB1:URA3/P<sub>TUB1</sub>GFP-TUB1:URA3, Spc105-FRB:KanMX/Spc105-FRB:KanMX, P<sub>Rec8</sub>-Spc105<sup>RASA</sup>:LEU2/P<sub>Rec8</sub>-Spc105<sup>RASA</sup>:LEU2, mek1::KanMX/mek1::KanMX |
| LY6995 | MATa/$\alpha$, tor1-1::HIS3/tor1-1::HIS3, fpr1::NatMX/fpr1::natMX4, RPL13A-2XFKBP12:TRP1/RPL13A-2XFKBP12:TRP1, Spc42-mCherry:HphMX/+, P<sub>TUB1</sub>GFP-TUB1:URA3/P<sub>TUB1</sub>GFP-TUB1:URA3, Spc105-FRB:KanMX/Spc105-FRB:KanMX, P<sub>Rec8</sub>-Spc105<sup>RASA</sup>:LEU2/P<sub>Rec8</sub>-Spc105<sup>RASA</sup>:LEU2, mad3::KanMX/mad3::KanMX |
| LY2606 | MATa/$\alpha$ mek1::KanMX6/ mek1::KanMX6, Spc42-mCherry:HphMX/+, P<sub>TUB1</sub>GFP-TUB1:LEU2/ P<sub>TUB1</sub>GFP-TUB1:LEU2, ZIP1-GFP/+ |
| LY10111 | MATa/$\alpha$, tor1-1::HIS3/tor1-1::HIS3, fpr1::NatMX/fpr1::natMX4, RPL13A-2XFKBP12:TRP1/RPL13A-2XFKBP12:TRP1, Spc105-FRB:KanMX/Spc105-FRB:KanMX, P<sub>Rec8</sub>-Spc105<sup>RASA</sup>:LEU2/P<sub>Rec8</sub>-Spc105<sup>RASA</sup>:LEU2, Bub3-eGFP:TRP1/Bub3-eGFP:TRP1, Spc42-mCherry:HphMX/+, mek1::HphMX/mek1::HphMX |
| LY9592 | MATa/$\alpha$, tor1-1::HIS3/tor1-1::HIS3, fpr1::NatMX/fpr1::natMX4, RPL13A-2XFKBP12::loxP/RPL13A-2XFKBP12::loxP, Spc105-FRB:KanMX/Spc105-FRB:KanMX, P<sub>Rec8</sub>-Spc105<sup>RASA</sup>:LEU2/P<sub>Rec8</sub>-Spc105<sup>RASA</sup>:LEU2, Bub3-eGFP:TRP1/Bub3-eGFP:TRP1, Spc42-mCherry:HphMX/+ |
| LY9946 | MATa/$\alpha$, Spc42-mCherry:HphMX /+, P<sub>CUP1</sub>- GFP-Rec8-LacI:HIS3/ P<sub>CUP1</sub>- GFP-Rec8-LacI:HIS3, LacO:TRP1 at 800 kb from CEN4/ LacO:TRP1 at 800 kb from CEN4, mek1::KanMX/mek1::KanMX, *spc105*<sup>RVAF</sup>/*spc105*<sup>RVAF</sup> |
| LY10032 | MATa/$\alpha$, Spc42-mCherry:HphMX /+, P<sub>CUP1</sub>- GFP-Scc1-LacI:HIS3/ P<sub>CUP1</sub>- GFP-Scc1-LacI:HIS3, LacO:TRP1/ LacO:TRP1, Spc42-mCherry:HphMX/+, cdc6::HisG/cdc6::HisG, P<sub>Gal1</sub>-ubiCdc6:URA3/ P<sub>Gal1</sub>-ubiCdc6:URA3, *spc105*<sup>RVAF</sup>/*spc105*<sup>RVAF</sup> |
| LY10092 | MATa/$\alpha$, Spc42-mCherry:HphMX /+, P<sub>CUP1</sub>- GFP-Scc1-LacI:HIS3/ P<sub>CUP1</sub>- GFP-Scc1-LacI:HIS3, LacO:TRP1 at 800 kb from CEN4/ LacO:TRP1 at 800 kb from CEN4, bub3::Leu2/bub3::Leu2, trp1::Bub3-3mCherry:TRP1/trp1::Bub3-3mCherry:TRP1, cdc6::HisG/cdc6::HisG, P<sub>Gal1</sub>-ubiCdc6:URA3/ P<sub>Gal1</sub>-ubiCdc6:URA3, *spc105*<sup>RVAF</sup>/*spc105*<sup>RVAF</sup> |
| LY9351 | MATa/$\alpha$, tor1-1/tor1-1::HIS3, fpr1::NatMX/fpr1::natMX4, RPL13A-2XFKBP12:TRP1/RPL13A-2XFKBP12::loxP, Ipl1-FRB:KanMX/Ipl1-FRB:KanMX, Spc42-mCherry:HphMX/+, mek1::HphMX/mek1::HIS3MX |
| LY10112 | MAT a/$\alpha$, tor1-1/tor1-1, fpr1::NatMX/fpr1::natMX4, RPL13A-2XFKBP12::loxP/RPL13A-2XFKBP12::loxP, Mps1-FRB:KanMX/Mps1-FRB:KanMX, Spc42-mCherry:HphMX/+, mek1::HIS3MX/mek1::HIS3MX |
| LY10113 | MAT a/$\alpha$, Ipl1-3GFP:NatMX/Ipl1-3GFP:NatMX, Mtw1-mRuby2:His5/Mtw1-mRuby2:His5 |
| LY10114 | MAT a/$\alpha$, Ipl1-3GFP:NatMX/Ipl1-3GFP:NatMX, Mtw1-mRuby2:His5/Mtw1-mRuby2:His5, mek1::KanMX/mek1::KanMX |
| LY5857 | MAT a/$\alpha$, Bub3-eGFP:TRP1/Bub3-eGFP:TRP1, Spc42-mCherry:hph/+ |
| LY9670 | MAT a/$\alpha$, Bub3-eGFP:TRP1/Bub3-eGFP:TRP1, mCherry-Tub1:ADE2/mCherry-Tub1:ADE2, mCherry-Tub1:URA3/mCherry-Tub1:URA3 |
| LY9669 | MAT a/$\alpha$, Bub3-eGFP:TRP1/Bub3-eGFP:TRP1, mCherry-Tub1:ADE2/mCherry- |
|  | Tub1::ADE2, mCherry-Tub1::URA3/mCherry-Tub1::URA3,<br>mek1::NatMX/mek1::NatMX |
| LY9961 | MATa/$\alpha$, Spc42-mCherry:HphMX/+, Mad2-3GFP:KanMX/Mad2-3GFP:KanMX,<br>mek1::HIS3MX/mek1::HIS3MX |
| LY5791 | MATa/$\alpha$, Spc42-mCherry:HphMX/+, Mad2-3GFP:KanMX/Mad2-3GFP:KanMX |
| LY8397 | MATa/$\alpha$, Spc42-mCherry:HphMX/+ Rec8-yEGFP:KanMX/ Rec8-yEGFP:KanMX,<br>spo12::HIS3MX/spo12::HIS3MX |
| LY8473 | MATa/$\alpha$, Spc42-mCherry:HphMX/+, Rec8-yEGFP:KanMX/ Rec8-yEGFP:KanMX,<br>spo12::HIS3MX/spo12::HIS3MX, mad3::KanMX/mad3::KanMX |
| LY10046 | MATa/$\alpha$, Spc42-mCherry:HphMX/+, Rec8-yEGFP:KanMX/ Rec8-yEGFP:KanMX,<br>spo12::HIS3MX/spo12::HIS3MX, ama1::NatMX/ama1::NatMX |
| LY9322 | MATa/$\alpha$, Spc42-mCherry:HphMX/+, Rec8-yEGFP:KanMX/ Rec8-yEGFP:KanMX |
| LY8599 | MATa/$\alpha$, Spc42-mCherry:HphMX/+, Rec8-yEGFP:KanMX/ Rec8-yEGFP:KanMX,<br>bub3::Leu2/bub3::Leu2, trp1::Bub3-3mCherry:TRP1/ trp1::Bub3-3mCherry:TRP1 |
| LY8598 | MATa/$\alpha$, Spc42-mCherry:HphMX/+, Rec8-yEGFP:KanMX/ Rec8-yEGFP:KanMX,<br>bub3::Leu2/bub3::Leu2, trp1::Bub3-3mCherry:TRP1/ trp1::Bub3-3mCherry:TRP1,<br>spo12::HIS3MX/spo12::HIS3MX |
| LY10047 | MATa/$\alpha$, Spc42-mCherry:HphMX/+, Rec8-yEGFP:KanMX/ Rec8-yEGFP:KanMX,<br>bub3::Leu2/bub3::Leu2, trp1::Bub3-3mCherry:TRP1/ trp1::Bub3-3mCherry:TRP1,<br>spo12::HIS3MX/spo12::HIS3MX, ama1::HphMX/ama1::HphMX |

**Table S2.** Plasmids used in this study.

| Plasmid name | Relevant subcloned regions | Purpose | Source |
| --- | --- | --- | --- |
| pLB304 | P<sub>Cup1</sub>-GFP-Scc1-LacI-mCherry:His | Mitosis separase biosensor | Yaakov et al., 2012 |
| pLB305 | P<sub>Cup1</sub>-GFP-Rec8-LacI:His | Meiosis separase biosensor | Yaakov et al., 2012 |
| pSLB22 | LacO:Trp1 | LacO repeats at the *TRP1* locus | Straight et al., 1996 |
| pLB113 | P<sub>CUP1</sub>-LacI-GFP:His3 | LacI-GFP under the *CUP1* promoter at the *HIS3* locus | Straight et al., 1996 |
| pLB358 | P<sub>SPC105</sub>-SPC105<sup>WT</sup>:Leu2 | Spc105 under its endogenous promoter at the *LEU2* locus | This study |
| pLB300 | P<sub>Rec8</sub>-SPC105<sup>WT</sup> at Trp1 | Spc105 under the Rec8 promoter at the *TRP1* locus | This study |
| pLB321 | P<sub>Rec8</sub>-SPC105<sup>WT</sup> at Leu2 | Spc105 under the Rec8 promoter at the *LEU2* locus | This study |
| pLB365 | P<sub>SPC105</sub>-SPC105<sup>RASA</sup> at Leu2 | *spc105*<sup>RASA</sup> under its endogenous promoter at the *LEU2* locus | This study |
| pLB302 | P<sub>Rec8</sub>-SPC105<sup>RASA</sup> at Trp1 | *spc105*<sup>RASA</sup> under the Rec8 promoter at the *TRP1* locus | This study |
| pLB318 | P<sub>Rec8</sub>-SPC105<sup>RASA</sup> at Leu2 | *spc105*<sup>RASA</sup> under the Rec8 promoter at the *LEU2* locus | This study |

**Table S3.** List of reagents used in this study.

| Reagent | Source | Identifier |
| --- | --- | --- |
| Yeast extract | Thermo Fisher<br>Scientific | Ref#212720, Lot#2352585 |
| Peptone | Thermo Fisher<br>Scientific | Ref#211820, Lot#1263496 |
| Dextrose (D-Glucose)<br>Anhydrous | Fisher Chemical | Lot#204841 |
| Potassium acetate | Fisher Bioreagents | Lot#210889 |
| Yeast nitrogen base<br>without amino acids | Thermo Fisher<br>Scientific | Ref#291920, Lot#9148845 |
| Synthetic complete<br>mixture drop-out:<br>Complete | Formedium | Ref# DSCK2500, Batch# FM0A416/006650 |
| Bacto-agar | Thermo Fisher<br>Scientific | Cas#: 214010 |
| β-estradiol | Sigma | Cas#: 50-28-2 |
| Rapamycin | Fisher BioReagents | Cas#: 53123-88-9 |
| Alpha factor | Zymo research | Cas#: Y1001 |
| Nocodazole | Sigma | Cat# M1404 |
| Concanavalin A | Sigma | Cas#: 11028-71-0 |
| NaCl | Sigma | Cat# S5886-500G |
| Gentamicin | Sigma | Cat# G1272-10ML |
| Nourseothricin Sulfate<br>(Clonat) | Thermo Fisher | Cat# 50-103-5835 |
| Hygromycin B | Cornig | Product # 30-240-CR |
| Dimethyl sulfoxide<br>(DMSO) | Sigma | Cat#472301 |
| Ethanol | Fisher | Cas#64-17-5 |

## Notes

### Competing Interest Statement

The authors have declared no competing interest.

